# A global *Corynebacterium diphtheriae* genomic framework sheds light on current diphtheria reemergence

**DOI:** 10.1101/2023.02.20.529124

**Authors:** Melanie Hennart, Chiara Crestani, Sebastien Bridel, Nathalie Armatys, Sylvie Brémont, Annick Carmi-Leroy, Annie Landier, Virginie Passet, Laure Fonteneau, Sophie Vaux, Julie Toubiana, Edgar Badell, Sylvain Brisse

## Abstract

**Background:** Diphtheria, caused by *Corynebacterium diphtheriae*, reemerges in Europe since 2022. Genomic sequencing can inform on transmission routes and genotypes of concern, but currently, no standard approach exists to detect clinically important genomic features and to interpret emergence in the global *C. diphtheriae* population framework.

**Methods:** We developed the bioinformatics pipeline DIPHTOSCAN (available at https://gitlab.pasteur.fr/BEBP/diphtoscan) to extract from genomes of Corynebacteria of the *diphtheriae* species complex, medically relevant features including *tox* gene presence and disruption. We analyzed 101 human *C. diphtheriae* isolates collected in 2022 in metropolitan and overseas France (France-2022). To define the population background of this emergence, we sequenced 379 additional isolates (mainly from France, 2018-2021) and collated 870 publicly-available genomes.

**Results:** The France-2022 isolates comprised 45 *tox*-positive (44 toxigenic) isolates, mostly imported, belonging to 10 sublineages (<500 distinct core genes). The global dataset comprised 245 sublineages and 33.9% *tox-* positive genomes, with DIPHTOSCAN predicting non-toxigenicity in 16.0% of these. 12% of the global isolates, and 43.6% of France-2022 ones, were multidrug resistant. Convergence of toxigenicity with penicillin and erythromycin resistance was observed in 2 isolates from France-2022. Phylogenetic lineages Gravis and Mitis contrasted strikingly in their pathogenicity-associated genes.

**Conclusions:** This work provides a bioinformatics tool and global population framework to analyze *C. diphtheriae* genomes, revealing important heterogeneities in virulence and resistance features. Emerging genotypes combining toxigenicity and first-line antimicrobial resistance represent novel threats. Genomic epidemiology studies of *C. diphtheriae* should be intensified globally to improve understanding of reemergence and spatial spread.

## Introduction

Diphtheria was a leading cause of infant mortality before the implementation of anti-toxin therapy and mass vaccination programs. Classical diphtheria is a respiratory infection mainly caused by the *tox* gene-positive strains of the bacterium *Corynebacterium diphtheriae.* The disease is classically characterized by the presence of pseudomembranes on the tonsils, pharynx and larynx. Only some strains of *C. diphtheriae* can produce the diphtheria toxin, which is encoded by the *tox* gene carried by a prophage integrated into the chromosome of these strains. The toxigenic strains can induce severe systemic symptoms that include myocarditis and peripheral neuropathies. Other forms of infection include bacteriemic infections, most often caused by non-toxigenic strains, and cutaneous infections, which are considered to play an important role in the transmission of the pathogen.

Diphtheria has been virtually eliminated by mass vaccination, but can cause large outbreaks where vaccination coverage is insufficient (du Plessis et al., 2017; Polonsky et al., 2021; Badell et al., 2021). In France, no case was reported between 1990 and 2001 (Bonmarin et al., 2009), and in the 2017-2021 period only 6.4 *tox*-positive *C. diphtheriae* were detected per year by the French surveillance (our unpublished data). In striking contrast, in 2022, 45 *tox*-positive isolates were detected, including 34 from metropolitan France, mostly associated with recent arrival from abroad. *C. diphtheriae* also reemerges in several European countries, strongly associated with non-vaccinated young adults with cutaneous infections with a travel history from Afghanistan and other countries (Badenschier et al., 2022; Kofler et al., 2022).

Whole genome sequencing (WGS) is a powerful approach to understand transmission and define the pathogenicity-associated characteristics of infectious isolates. *C. diphtheriae* is a genetically diverse species with multiple phylogenetic sublineages among which a large heterogeneity of virulence or antimicrobial resistance factors is observed (Sangal & Hoskisson, 2016; Seth-Smith & Egli, 2019; Hennart et al., 2020; Guglielmini et al., 2021). One prominent polymorphism in *C. diphtheriae* is the variable presence of the *tox* gene, but the population dynamics and drivers of *tox* acquisition or loss remain poorly understood. In addition, non-toxigenic *tox*-bearing (NTTB) *C. diphtheriae* isolates represent 5-20% of *tox*-positive isolates, but our capacity to predict toxigenicity from genomic sequences is still limited. Several other experimentally-demonstrated virulence factors have been described in *C. diphtheriae* (Ott, 2018). Although early 1930s literature suggested a higher virulence of isolates of biovar Gravis (McLeod, 1943; Barksdale, 1970), it is unknown whether this historical observation applies to extant diphtheria cases, as recent Gravis isolates are more rarely *tox*-positive than those of biovar Mitis (Hennart et al., 2020). More generally, the population variation of virulence factors, and its interactions with clinical outcomes, remain largely to be characterized. Despite being rare, antimicrobial resistance (AMR) in *C. diphtheriae* is increasingly reported (Mina et al., 2011; Zasada, 2014; Forde et al., 2020; Hennart et al., 2020), but the mechanisms of resistance that are prevalent across world regions are not well known, and the evolutionary emergence and dissemination of multi-drug resistant *C. diphtheriae*, and its possible convergence with toxigenicity in the same strains, should be carefully monitored.

Although WGS of *C. diphtheriae* clinical isolates is increasingly performed for surveillance purposes, no simple tool currently exists for *C. diphtheriae* genomic feature extraction and interpretation in clinical, surveillance and research contexts. Besides, analyses of *C. diphtheriae* genomes remain largely unstandardized, which limits the interpretation of local genomic epidemiology studies in their global context. Advances towards standardization include the 7-gene MLST genotyping approach and attached nomenclature of sequence types (ST) (Bolt et al., 2010), and its core-genome MLST (cgMLST) extension and associated nomenclature of sublineages and genomic clusters (Guglielmini et al., 2021).

Here, we aimed to provide insights into the France 2022 diphtheria emergence by reporting on its epidemiology and by placing the involved isolates in the global genomic context of *C. diphtheriae* populations. We introduce DIPHTOSCAN, a genotyping tool designed for rapid and standardized genomic analyses of Corynebacteria of the *C. diphtheriae* species complex (CdSC), and illustrate its use by analyzing the 101 *C. diphtheriae* isolates (including 56 *tox*-negative ones) collected in 2022 in France (henceforth, the France-2022 dataset). We provide context of this emergence by analyzing 1249 other *C. diphtheriae* genomes of diverse geographic and temporal origins, including 379 newly sequenced isolates collected by the French national surveillance laboratory, mostly between 2018 and 2021. We uncovered novel insights into the global population structure of *C. diphtheriae*, including a striking contrast in pathogenesis-associated gene clusters between phylogenetic lineages Gravis and Mitis, and describe high-risk sublineages with convergence of resistance and virulence features.

## Material & Methods

### Clinical isolates inclusion and global genomic sequence dataset

To investigate the epidemiology of diphtheria in France, we included all cases of *C. diphtheriae* infections detected by the French surveillance in 2022. Among 144 isolates received by the National Reference Center, there were 101 deduplicated isolates when retaining only one from each patient. These were isolated in metropolitan France as well as in Mayotte, La Reunion and French Guiana (**France-2022 dataset, Table S1**). Note that metropolitan France comprises mainland France and Corsica, as well as nearby islands in the Atlantic Ocean, the English Channel (French: la Manche), and the Mediterranean Sea. All isolates collected in 2022 from metropolitan France were from mainland France. Overseas France is the collective name for all the French territories outside Europe.

In addition, a total of 1,249 comparative genomes were included (**Table S1**). First, we sequenced for the present study 379 additional isolates, including 320 collected prospectively between 2008 and 2021 by the French National Reference Center (NRC), 34 historical clinical isolates mostly from metropolitan France and 19 isolates from Algeria (Benamrouche et al., 2016). These new genomes were sequenced to complement the 226 previous genomes from *C. diphtheriae* from the French diphtheria surveillance system (Hennart et al., 2020; Guglielmini et al., 2021), including 43 isolates from Yemen (Badell et al., 2021). Together, these represent 599 produced by the NRC for Corynebacteria of the *diphtheriae* complex (**non-2022 French NRC dataset, Table S1**). Nearly four-fifths (531; 88.6%) of these isolates were prospectively collected between 2008 and 2021 from French metropolitan and overseas territories, 54 isolates (9.0%) were collected between 1990 and 2007 from France and Algeria and 14 (2.3%) isolates collected between 1951 and 1987 from metropolitan France.

Second, we included publicly-available genomes from NCBI, mostly previously published and isolated in South Africa (du Plessis et al., 2017), Germany-Switzerland (Meinel et al., 2016), Germany (Dangel et al., 2018; Berger et al., 2019), Canada (Chorlton et al., 2019), Austria (Schaeffer et al., 2020), the USA (Williams et al., 2020; Xiaoli et al., 2020), Spain (Hoefer et al., 2020), India (Will et al., 2021) and Australia (Timms et al., 2018). Altogether, this represents a dataset of 579 genomes (**non-French public dataset, Table S1**).

Further, we sequenced 6 ribotype reference strains (Grimont et al., 2004). Together with 65 previously sequenced (Hennart et al., 2020), this represents a dataset of 71 genomes of ribotype reference strains (**Table S1**).

From the global set of 1,249 genomes (**non-2022 French NRC + non-French public dataset + ribotype datasets**), we created a non-redundant subset of genomes by randomly selecting one genome per genomic cluster (threshold: 25 cgMLST mismatches; see below), isolation year and city (if city was unavailable, the country was used instead); this deduplicated subset comprised 976 genomes (hereafter, the ***global dataset***).

### Microbiological characterization of isolates at the French National Reference Laboratory

*C. diphtheriae* isolates were grown and purified on Tinsdale agar. Strains were characterized biochemically for pyrazinamidase, urease, and nitrate reductase and for utilization of maltose and trehalose using API Coryne strips (BioMérieux, Marcy l’Etoile, France) and the Rosco Diagnostica reagents (Eurobio, Les Ulis, France). The Hiss serum water test was used for glycogen fermentation. The biovar of isolates was determined based on the combination of nitrate reductase (positive in Mitis and Gravis, negative in Belfanti) and glycogen fermentation (positive in Gravis only). Antimicrobial susceptibility was determined by disc diffusion (BioRad, Marnes-la-Coquette, France). Zone diameter interpretation breakpoints are given in **Table S3**.

The presence of the diphtheria toxin *tox* gene was determined by real-time PCR assay (Badell et al., 2019), whereas the production of the toxin was assessed using the modified Elek test (Engler et al., 1997).

For genomic sequencing, isolates were retrieved from −80°C storage and plated on tryptose-casein soy agar for 24 to 48 h. A small amount of bacterial colony biomass was resuspended in a lysis solution (20 mM Tris-HCl [pH 8], 2 mM EDTA, 1.2% Triton X-100, and lysozyme [20 mg/ml]) and incubated at 37°C for 1 h. DNA was extracted with the DNeasy Blood&Tissue kit (Qiagen, Courtaboeuf, France) according to the manufacturer’s instructions. Genomic sequencing was performed using a NextSeq500 instrument (Illumina, San Diego, CA) with a 2 × 150-nucleotide (nt) paired-end protocol following Nextera XT library preparation (Hennart et al., 2020).

For de novo assembly, paired-end reads were clipped and trimmed using AlienTrimmer v0.4.0 (Criscuolo & Brisse, 2013), corrected using Musket v1.1 (Liu et al., 2013), and merged (if needed) using FLASH v1.2.11(Magoč & Salzberg, 2011). For each sample, the remaining processed reads were assembled and scaffolded using SPAdes v3.12.0 (Bankevich et al., 2012).

### Merging of the Oxford and Pasteur MLST databases

Two *C. diphtheriae* databases using the BIGSdb framework were originally designed separately for distinct purposes: while Oxford’s PubMLST database mainly offered 7-gene MLST (Bolt et al., 2010), the Pasteur database was used for the *Corynebacterium* cgMLST typing (Guglielmini et al., 2021). To facilitate the use of these resources and avoid redundancy in the curation of the two independent genomic libraries, a merging of the databases was decided in agreement with PubMLST administrators. In order to merge the data available in the two databases, we proceeded as per BIGSdb dual design: isolates genomes and provenance data were imported into the “isolates” database, whereas allelic definitions of MLST were imported into the “seqdef” database.

Regarding the isolates database, we first downloaded Oxford’s PubMLST *C. diphtheriae* database. To avoid isolate entries duplication, we identified common isolates between the two databases, and filtered duplicate isolates before import into the Pasteur database. In total, 684 out of 934 (73%) isolates from the Oxford database were imported. To facilitate the tracing of isolates and their possible previous existence in Oxford’s database, isolates identification numbers (BIGSdb-Pasteur ID number) of isolates from the Oxford database were numbered from 1,520 to 2,003. We also collated them into a public project collection called “Oxford” (project ID 13).

Regarding the sequence and profiles definition database, we imported MLST alleles and profiles into an initially void MLST scheme container within the BIGSdb-Pasteur database. MLST analysis was performed on all isolates of the BIGSdb-Pasteur database, including the ones imported from Oxford, which were therefore assigned the same MLST genotype as previously in the Oxford database.

At the end of the merging process, all isolates and MLST data from PubMLST’s *C. diphtheriae* database were available into the BIGSdb-Pasteur *C. diphtheriae* species complex database (https://bigsdb.pasteur.fr/diphtheria/), and Oxford’s PubMLST *C. diphtheriae* database was shut down. As of September 22^nd^, 2022, the database resulting from the merged datasets comprised 1,478 public isolates records with 794 associated genomes, and 2,392 isolates in total when considering private entries. The number of entries varied across species: *C. diphtheriae* (n = 1,291; 87.4%) and *C. ulcerans* (n = 131; 8.9%), *C. belfantii* (n = 45; 3.0%) and *C. rouxii* (n = 10; 0.7%). The MLST scheme comprised 854 registered STs.

### cgMLST and nomenclature of sublineages

The MLST and cgMLST genotypes (cgST) were defined using the Institut Pasteur *C. diphtheriae* species complex database at https://bigsdb.pasteur.fr/diphtheria.

A core genome MLST (cgMLST) scheme comprising 1,305 loci (Guglielmini et al., 2021) was employed to define the alleles and cgST of the 1,249 genomic sequences using BIGSdb (https://bigsdb.pasteur.fr/diphtheria). Using the 1,249-genomes dataset, the mean number of missing alleles per profile was 12 (0.9%) and almost all (n=1,242; 99.4%) genomes had a cgMLST profile with fewer than 65 (5%) missing alleles. A cgST number was defined for all but one cgMLST profiles (one genome had 219 missing alleles, whereas the admissible threshold is 10%, i.e., 130 missing alleles).

Genomes were classified using the single-linkage cluster-profile.pl function of BIGSdb into genomic clusters (25 mismatch threshold) and sublineages (500 mismatches). Sublineages were attributed numbers by using an ST inheritance rule (Hennart et al., 2022), which was applied from SL1 to SL744, after which the numbers are attributed consecutively with no reference to MLST identifiers, starting at 10,000 (see column ‘SL’ in **Table S1**).

### Phylogenetic analysis based on a core genome

Panaroo v1.2.3 was used to generate from the assembled genomic sequences, a core genome used to construct a multiple sequence alignment (cg-MSA). The genome sequences were first annotated using prokka v1.14.5 with default parameters, resulting in GFF files. Protein-coding gene clusters were defined with a threshold of 70% amino acid identity, and core genes were concatenated into a cg-MSA when present in 95% of genomes. IQtree version 2 was used to build a phylogenetic tree based on the cg-MSA, with the best fitting model TVM+F+R5. The tree was constructed from 1,948 core genome loci, for a total alignment length of 1,986,172 bp (79.8% of NCTC13129 genome length, of 2,488,635 bp), was rooted using *C. belfantii* strain FRC0043^T^, and is available at: https://itol.embl.de/tree/1579917435471751662784292.

### Development of the DIPHTOSCAN pipeline

To develop DIPHTOSCAN, we combined code from Kleborate v2.2.0 (Lam et al., 2021), AMRfinderPlus (Feldgarden et al., 2021) and BIGSdb (Jolley & Maiden, 2010) with some modifications. The structures of DIPHTOSCAN and its custom database are presented in **Figure S3** and **Figure S4**. A custom code was created for DIPHTOSCAN initiation, interpretation and for displaying results. The *C. diphtheriae* specific genes (genomic markers, AMR determinants and virulence factors) for which the genomes are screened by DIPHTOSCAN (**Figure S4**) are provided in a custom database similar in its structure to the AMRFinderPlus database (https://ftp.ncbi.nlm.nih.gov/pathogen/Antimicrobial_resistance/); this database can be further enriched with novel features in the future. When launching DIPHTOSCAN, the AMRFinderPlus and custom databases are merged. We used the functions of species determination, MLST genotyping, and full CDS prediction from Kleborate. All the functionalities are presented in **Figure S2**. To facilitate readability and downstream analyses, the output of DIPHTOSCAN is generated in a tab-delimited format. The execution time of DIPHTOSCAN increases linearly with the number of input genomes. Roughly, 40 seconds are needed to scan a single genome with 1 cpu. DIPHTOSCAN computations can be parallelized, as AMRFinderPlus and JolyTree use parallelization.

### Assignment of species, MLST and Sequence Types (ST)

To perform rapid and accurate species identification, DIPHTOSCAN uses the k-mer-derived Mash distances (Ondov et al., 2016). DIPHTOSCAN calculates Mash distances (Mash v2.2) between the query genomes and a collection of reference assemblies of the *CdSC*, and reports the species with the smallest distance. *C. diphtheriae* genomes were confirmed as *C. diphtheriae* based on a Mash distance smaller than 0.05 with either the *C. diphtheriae* type strain NCTC11397^T^ (= C7S), the reference genome strain NCTC13129, or the vaccine strain PW8 (Park-Williams 8).

Mash distance ≤0.05 is reported as a strong match, ≤0.1 as weak. We have used and adapted the structure of the Kleborate tool for this function. This approach was validated by comparing DIPHTOSCAN species assignments with those obtained by average nucleotide identity (ANI; Konstantinidis and Tiedje, 2005) using FastANI (Jain et al., 2018) using the global dataset; 100% concordance was achieved.

MLST profiles and sequence types (ST) were defined using the international MLST scheme for *C. diphtheriae* and *C. ulcerans*. DIPHTOSCAN defines these genotypes for genomic sequences using the analogous script from Kleborate. In order to use an up-to-date version of the MLST nomenclature, which is regularly updated, the MLST profiles and alleles are downloaded at the start of the pipeline before genotyping the genomes. The download_alleles.py script from BIGSdb is used for this purpose (https://github.com/kjolley/BIGSdb/tree/develop/scripts/rest_examples).

### Biovar-associated markers detection

The three main biovars of *C. diphtheriae* can be distinguished based on isolate abilities to reduce nitrate and to metabolize glycogen. Previously, a strong concordance was found between the biovar and the presence in the genome of several genomic markers including *spuA*, which codes for a putative alpha-1,6-glycosidase, and the *narKGHJI* operon for nitrate reductase (Sangal et al., 2014; Santos et al., 2018; Hennart et al., 2020). We therefore included in the custom DIPHTOSCAN query database the *spuA* marker and its adjacent genes (DIP0351; DIP0353; DIP0354; DIP0357=*spuA*), which are strongly associated with biovar Gravis, and the *narIJHGK* cluster, which is typically absent or partly disrupted, mainly due to mutations in the *narG* (Hennart et al., 2020) or *narI* (Sangal et al., 2014) in isolates of biovar Belfanti. In the future, markers of the two biovars of *C. pseudotuberculosis* may be added.

### Detection of antibiotic resistance genes

Antibiotic resistant genes were identified using AMRfinderPlus, with the database found at: https://ftp.ncbi.nlm.nih.gov/pathogen/Antimicrobial_resistance/. Features are detected by using the BLAST family of tools, with identity and coverage defined for each family of antibiotics (fam.tab). A few genes particularly relevant for the *CdSC* were added to this database: *pbp2m* (Forde et al., 2020; Hennart et al., 2020) and mutation points of *rpoB* (WP_004566675.1) and *gyrA* (WP_010933942.1). AMRfinderPlus v3.11.2 is used within DIPHTOSCAN with no modifications.

### Detection of virulence genes from the *C. diphtheriae* species complex

A custom database of virulence features of *C. diphtheriae* and related species was compiled from literature for the purposes of this work. We included in the custom query database, a panel of genetic features for which published experimental evidence of their clinical relevance exists in *C. diphtheriae* or closely related species (*i.e.,* increased virulence in animal models, or decreased antimicrobial susceptibility *in vitro*) (**Table S2**). These target genes are the following: *tox*, SpaA-, SpaD-, and SpaH-type pili gene clusters, DIP0733 (*67-72p*), the genes DIP1281 and DIP1621 that code for proteins of the NlpC/P60 family, DIP0543 (*nanH*), DIP1546 and DIP2093 (Ott, 2018) and *pld* (phospholipase). A second panel of genetic features with no experimental evidence but with strong suspicion for a role in virulence, based on homology with genes from other pathogens, was also included for broader screening of virulence features (**Table S2**).

For the main virulence factor, the *tox* gene, we used a reference sequence of this gene from each of *C. diphtheriae*, *C. ulcerans* and *C. pseudotuberculosis* (WP_003850266.1, WP_014835773.1 and WP_014654963.1, respectively), as the toxin differs between these species (Dangel et al., 2019).

The *tox* gene may be disrupted in some strains by the occurrence of stop codons or other genetic events, leading to non-toxigenic, *tox*-gene bearing (NTTB) isolates (Zakikhany et al., 2014; Melnikov et al., 2022). DIPHTOSCAN provides information on the putative toxicity of a strain from the *tox* gene sequence using a categorization into four possible outputs, following the convention proposed in Kleborate (Lam et al., 2021): (i) if the sequence in the analyzed genome is identical to the reference tox sequence from NCTC13129 strain, the output provides the name of the sequence with the denomination of the species (*e.g.,* tox_diphtheriae); (ii) If the sequence in the analyzed genome has a coverage length identical to the reference, but an identity different from 100%, then an asterisk (*) is added (*e.g.,* tox_diphtheriae*); (iii) If the hit coverage length is smaller than the reference length, the tag ‘-NTTB?-xx%’ is added, where xx is the percentage of the missing sequence length compared to the reference length); (iv) Finally, if the truncated *tox* sequence is located at the end of a contig, the symbol ‘$’ is added, to highlight that the prediction is uncertain.

To analyse the *tox* gene promotor region, the sequence of strain NCTC13129 corresponding to the 300 nt upstream of *tox* start codon was used as a query in BLASTn analyses. Sequence alignment of the corresponding region in the queried genomes was performed with seaView and the mutations were visualized and compared with the distribution of the available results of the Elek test. For DtxR sequence variation, *dtxR* was detected using BLASTn with DIPHTOSCAN, and the translation into amino acid, alignment and visualization of mutations were performed using seaView.

Virulence genes were identified using the method of AMRfinderPlus but based on our custom database of virulence features. The virulence genes are detected by BLASTn with thresholds of minimum 80% identity and 50% coverage. Based on the output of AMRfinderPlus, the gene completion and allele similarity is reported as described above for the *tox* gene following the Kleborate convention.

## Results

### 1. The re-emergence of *C. diphtheriae* in France in 2022

In 2022, the French NRC has received 101 human samples of *C. diphtheriae,* from metropolitan France (n=76) as well as in the Indian Ocean islands of Mayotte (n=10) and La Reunion (n=6), and in French Guiana (n=9). There were 45 isolates carrying the *tox* gene coding for diphtheria toxin (*tox*-positive isolates), whereas in the five previous years a total of 32 *tox*-positive *C. diphtheriae* were detected (**Figure S1A**). *C. diphtheriae* were isolated in metropolitan France (n=34) and in Mayotte/La Reunion (n=11), while none were found in French Guiana. The metropolitan France isolates were isolated only in the second part of the year (**Figure S1B**) and were associated with a recent travel history from Afghanistan (n=24) or other countries from West Africa, North Africa, Middle East and Southern Asia; These isolates were predominantly from cutaneous infections, whereas 7 were from respiratory infections (**Table S1**; **Figure 1**). Only 3 of the 34 patients were up to date with their vaccination.

**Figure 1.**
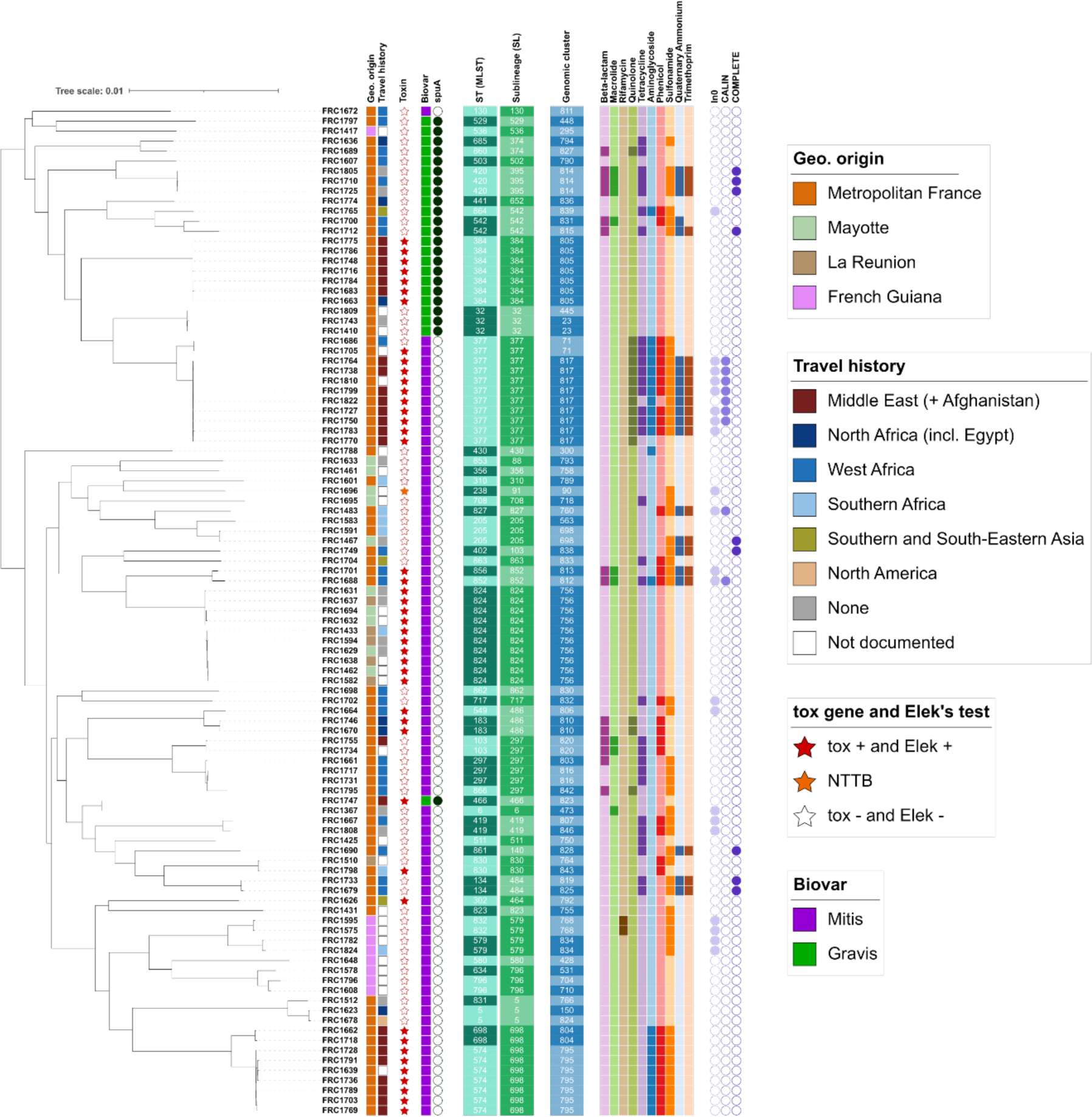
Phylogenetic tree of *Corynebacterium diphtheriae* from France, 2022. The tree was obtained by maximum likelihood based on a multiple sequence alignment of the core genome. The scale bar represents the number of nucleotide substitutions per site. The first column that follows the isolates identifiers indicates the geographic origin (place of isolation; see key). Travel history provides the most distant geographic region of reported travel (see key); note that Afghanistan was included in Near and Middle East; and Egypt was included in North Africa. The stars represent the presence (red star), presence but disruption (NTTB, orange) or absence (white star) of the diphtheria toxin tox gene. Biovars are represent in colored squares, and spuA gene presence by a dark green circle. MLST STs, sublineage (SL) and genomic clusters are provided with an alternation of colored strips. Identifiers of the main STs are indicated (note the strong concordance between ST and cgMLST sublineages). The 10 next colored columns correspond to the presence of at least one gene or mutation (for quinolone and rifamycin classes) involved in resistance to the indicated class of antimicrobial agents. Last, the presence of integron-related structures *(Cury et al., 2016)* is indicated: In0 (integron integrase and no attC sites), CALIN (clusters of attC sites lacking integron-integrases) and complete integrons (integrase and at least one attC site). The simultaneous presence of In0 and CALIN may denote their presence in different contigs even though the integron might be complete.

### 2. Development of the DIPHTOSCAN pipeline

To provide a tool to extract information from genomes of *C. diphtheriae* and related potentially toxigenic species, we developed DIPHTOSCAN. The technical characteristics of DIPHTOSCAN are summarized in **Figure S2-S4** and the methodological details for genotyping are provided in the Methods section.

In brief, the DIPHTOSCAN pipeline (**Figure S2**) starts with taxonomic assignment of species. Recent taxonomic updates have defined, besides the three classical species *C. diphtheriae*, *C. ulcerans* and *C. pseudotuberculosis*, three novel species of the Corynebacteria of the *diphtheriae* species complex (CdSC): *C. belfantii* (Dazas et al., 2018)*, C. rouxii* (Badell et al., 2020) and *C. silvaticum* (Dangel et al., 2020). If the genome is confirmed to belong to the CdSC, 7-gene MLST analysis (Bolt et al., 2010) is performed. For *C. diphtheriae*, additional genotype categorizations can be performed using the BIGSdb-Pasteur database tool: cgST, genomic cluster and sublineage assignment (Guglielmini et al., 2021). Next, the detection of antimicrobial resistance determinants (mutations in core genes and horizontally acquired genes) and virulence factors is performed. DIPHTOSCAN also includes a prediction of the functionality or disruption of the *tox* gene, the most important virulence factor of CdSC isolates. DIPHTOSCAN next searches for genomic markers associated with biovars Gravis, Mitis and Belfanti, a biochemical-based classification that was initiated in the 1930s (Anderson et al., 1931; McLeod, 1943) and which is still in use for *C. diphtheriae* strain characterization. IntegronFinder2 (Néron et al., 2022) was included in the pipeline to contextualize resistance genes. Last, a rapid phylogenetic method based on k-mer distances, JolyTree (Criscuolo, 2020), was integrated to provide quick phylogenetic trees for the genomic assembly datasets under study. The two latter steps are optional.

### 3. Genetic diversity of *C. diphtheriae* isolates from France, 2022

The *C. diphtheriae* isolates belonging to the France-2022 dataset were sequenced and their genomic sequences were analyzed using DIPHTOSCAN. Sublineage classification of the isolates showed that the France-2022 dataset comprised 41 distinct sublineages (defined using the 500 cgMLST mismatch threshold). The nomenclature of these sublineages was established using an inheritance rule that captures their majority MLST denomination, where possible (Guglielmini et al., 2021; Hennart et al., 2022), resulting in a strong concordance of sublineage denominations with the classical MLST identifiers (**Figure 1**). There were 51 different STs, as 9 sublineages comprised two or more closely related STs; in 7 of 9 cases, they only differed by a single locus. Sublineages thus appeared as useful classifiers for closely related STs.

There were four frequently isolated *tox*-positive sublineages: SL824 included 10 isolates from Mayotte and La Reunion; these all belonged to the same genomic cluster (GC756), indicating recent transmission. Three other frequent *tox*-positive sublineages were SL377 (n=11 isolates, 10 of which were *tox*-positive), SL698 (n=9) and SL384 (n=7), which were associated with travel from Afghanistan and countries of the Middle East (**Figure 1**). Whereas SL384 was genetically homogeneous (GC805), SL377 and SL698 both comprised two genomic clusters (SL377: GC817 and GC71; SL698: GC795-ST574 and GC804-ST698). SL377-GC71 was not associated with Afghanistan and one isolate from Senegal was *tox*-negative.

Besides the above four frequent sublineages, six additional *tox*-positive sublineages were isolated: three isolates of sublineage SL486 associated with Senegal and Tunisia; two SL852 isolates associated with Mali; and one SL466 isolate associated with travel from Afghanistan and one SL464 isolate associated with Thailand. SL91 comprised one non-toxigenic, *tox*-gene bearing (NTTB) isolate, and SL830 comprised 2 isolates: one *tox*-positive and one *tox*-negative.

Besides, there were 31 *tox*-negative sublineages, which were typically isolated once or twice only; a notable exception was SL297, which comprised six *tox*-negative isolates associated with travel from Egypt, Senegal, and Mali (**Figure 1**).

### 4. The global phylogenetic framework of *C. diphtheriae*

We investigated the global diversity of *C. diphtheriae* to provide context to the France-2022 emerging genotypes. A dataset of 1,249 comparative *C. diphtheriae* genomes were sequenced or gathered from previous studies (see Methods). cgMLST grouped these isolates into 245 sublineages. The 7-gene MLST analysis revealed 364 distinct STs. Almost all (360; 98.6%) STs corresponded one-to-one with the sublineage level, *i.e*., all isolates of these STs belonged to the same sublineage. However, 72 sublineages (29.4%) comprised at least two STs. Of the 123 novel sublineages uncovered here, 114 sublineages were given an identifier inherited from the 7-gene MLST nomenclature (whereas 9 were attributed an arbitrary number, see Methods).

There were 576 genomic clusters, many of which comprised previously documented epidemiological clusters of related isolates. For example, GC456 comprised 43 isolates from a Vancouver inner city outbreak (Chorlton et al., 2019). Whereas 47 GCs had between 5 and 27 isolates (**Table S1**; **Figure S5A**), the 529 remaining ones had only 1 and 4 isolates. 106 (43.3%) of the 245 sublineages comprised at least two genomic clusters.

To eliminate the population bias introduced by multiple sampling of outbreak strains, we created a non-redundant subset by randomly selecting one genome per genomic cluster, isolation year and city (if city was unavailable, the country was used instead) and with the same resistance genes profile and *tox* status (see column ‘Dataset’ in **Table S1**). These 976 deduplicated genomes (hereafter, the *global dataset*) define the background population of *C. diphtheriae*.

Within the global dataset, 35 sublineages were represented 7 times of more (**Figure 2**). The two predominant sublineages were SL8 (n=61) and SL5 (n=48); their main 7-gene MLST sequence types were ST8 and ST5, previously noted to be predominant in the ex-USSR 1990s outbreak. The most represented *tox*-positive sublineages in the global dataset were SL8, SL453, SL486, SL377 and SL91, and SL50 was a predominant NTTB sublineage (**Figure 2**).

**Figure 2.**
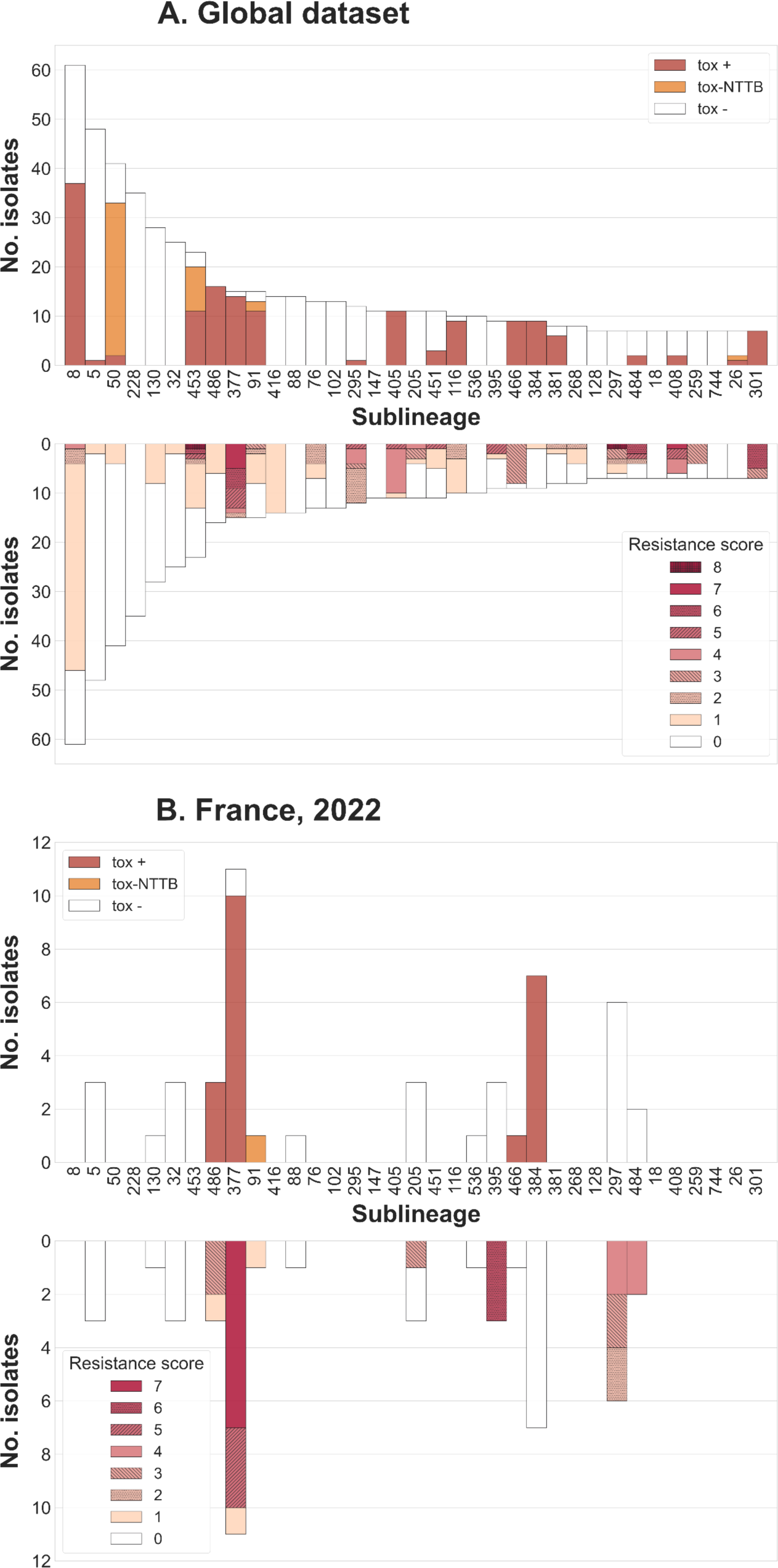
Sublineage distribution of tox gene and resistance score. (Top) Bar length correspond to the number of isolates per sublineage (deduplicated global dataset, 976 isolates). Upper part: isolates with non-disrupted tox are colored in red, with disrupted tox (NTTB) in orange, and not carrying the tox gene in white. Lower part: bar sectors are colored by resistance score (including beta-lactams and macrolides; see key). (Bottom) Bar length correspond to the number of isolates per sublineage (France, 2022 dataset, 101 isolates). Bar sectors are colored as in the top panel.

Of the 10 sublineages with *tox*-positive isolates observed in France-2022, 7 were found in the global dataset; of which 5 were among the 35 frequent global sublineages. Besides, 9 *tox*-negative sublineages from France-2022 were also frequent in the global dataset (**Figure 2**). Of the common France-2022 sublineages, SL377, SL384 and SL297 were also common in the global dataset (**Figure 2**), and their toxigenicity and resistance features matched those observed in the global dataset. In contrast, SL698 (metropolitan France) and SL824 (Indian Ocean) were uniquely common in the France-2022 dataset (**Figure S5B**).

The phylogenetic structure of *C. diphtheriae* revealed a star-like phylogeny with multiple deeply-branching sublineages as previously reported (Berger et al., 2019; Seth-Smith & Egli, 2019; Hennart et al., 2020; Guglielmini et al., 2021) (**Figure 3**). Sublineages appeared to be grouped according to biovars Gravis (and its *spuA* marker gene) and Mitis as previously noted (Hennart et al., 2020), as they formed two main lineages named Gravis (green branches) and Mitis (purple), defined by the presence of the *spuA* gene (**Table S1**). cgMLST-defined sublineages were highly concordant with the phylogeny and often comprised more than one 7-gene ST (**Figure 3; Table S1**). The frequent *tox*-positive sublineages SL377 and SL384 were phylogenetically related within lineage Gravis (**Figure 3**), suggesting they share ancestrally-acquired genetic features.

**Figure 3.**
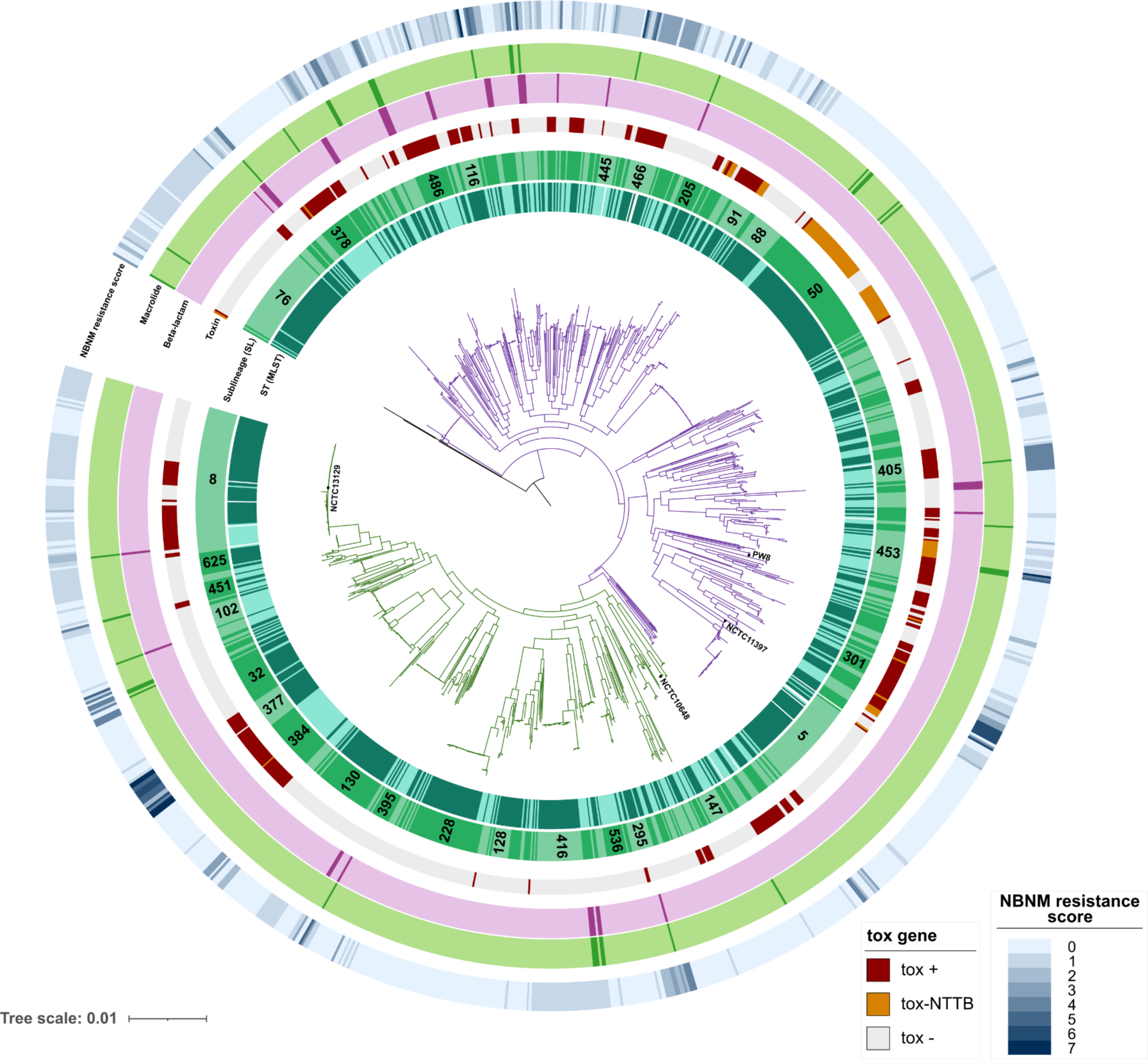
Phylogenetic tree of *Corynebacterium diphtheriae*. The tree was obtained by maximum likelihood based on a multiple sequence alignment of the core genome, and was rooted with C. belfantii (not shown). The scale bar gives the number of nucleotide substitutions per site. The main lineages Mitis and Gravis are drawn using purple and green branches, respectively. The two inner circles indicate MLST and sublineage alternation, respectively; main sublineages are labeled within the sectors. first ten colored circles around the tree correspond to the different classes of antibiotics. The following circle indicates the presence, disruption or absence of the diphtheria toxin tox gene (see key). The beta-lactam resistance circle indicates the presence of the pbp2m gene, while the macrolide circle corresponds to the presence of *ermX* or *ermC* (darker color: presence of the genomic determinant). The most external circle indicates the non-beta-lactam, non-macrolide (NBNM) resistance score (number of classes with at least one resistance feature), as a blue gradient (see key). Four reference strains are indicated: strain NCTC13129, which is used as genomic sequence reference; strain NCTC10648, which is used as the tox-positive and toxinogenic reference strain in PCR and Elek tests, respectively; strain NCTC11397^T^, which is the taxonomic type strain of the C. diphtheriae species; and the vaccine production strain PW8.

We placed within this population background, the France-2022 isolates (**Figure S6**), which appeared to be dispersed in multiple branches of the global phylogeny. The isolates previously collected by the French reference laboratory appeared even more diverse and largely dispersed across the global phylogenetic diversity of *C. diphtheriae* (**Figure S6**), indicating that a large fraction of the global diversity has been sampled by the French surveillance system.

Ribotyping was previously used as a classification and nomenclature system of *C. diphtheriae* strains (Grimont et al., 2004; Mokrousov, 2009). The 71 ribotype reference strains sequenced herein or previously (Hennart et al., 2020) were placed in the global phylogeny (**Figure S7**), showing that these strains are highly diverse. However, this ribotype subset is biased towards tox-positives (40 of 71 strains) and appears to represent unevenly and incompletely, the currently sampled *C. diphtheriae* diversity.

### 5. Population distribution of the diphtheria toxin gene

To evaluate DIPHTOSCAN for its ability to detect the *tox* gene and to predict its toxigenicity, we used the 855 isolates for which data on *tox* qPCR and Elek test were available. DIPHTOSCAN detected that *tox* was located at the end of a contig and therefore incomplete in 3 cases (reported with a ‘$’ suffix, indicating genomic assembly truncation). Of the 852 remaining isolates, 221 were *tox*-positive and 631 *tox*-negative by the reference qPCR method. DIPHTOSCAN detected the *tox* gene in 219 (99.1%) of the *tox*-positives, and reported its absence in 2 isolates. Among the 631 *tox*-negative isolates, DIPHTOSCAN reported the absence of the gene in 625 (99.0) isolates. Of 198 Elek-positives, 195 (98.5%) were predicted to be toxigenic by DIPHTOSCAN, whereas 1 was predicted to be non-toxigenic and for two isolates the *tox* gene was not detected. Of the Elek-negative isolates, 11 (50.0%) were predicted as non-toxigenic by DIPHTOSCAN. Thus, *tox* detection by DIPHTOSCAN was both sensitive and specific, whereas toxigenicity prediction was highly sensitive but not highly specific, likely due to unexplained non-toxigenicity in isolates with a full-length toxin gene. However, we did not observe non-toxigenicity-associated variation in the promoter region of the *tox* gene, nor on the DtxR protein sequence.

In the France 2022 dataset, 45 genomes were detected as *tox*-positive and 44 of these were predicted as toxigenic, with 100% concordance with the Elek test. In comparison, within the global dataset, approximately one third of the isolates (331/976; 33.9%) were *tox*-positive, as defined using DIPHTOSCAN, which detected a truncation and hence predicted non-toxigenicity in 16.0% of these (52/331).

Combining the global dataset with the France 2022 dataset (1077 genomes), DIPHTOSCAN identified 33 *tox* alleles. Among these, the most frequent are *tox*-2 (n=97, including the vaccine strain PW8), *tox*-3 (n=81) and *tox*-1 (n=41). These alleles are synonymous and thus result in the same amino acid sequence of the diphtheria toxin, implying complete match with the vaccine strain toxin. Alleles *tox*-24, 25, 26,27, 35, 36, 37 and 38 were predicted as NTTB by DIPHTOSCAN. The potential impact of protein changes deduced from *tox* gene sequence variation was previously analyzed (Will et al., 2021); we provide the correspondence of *tox* alleles in this previous study and ours in **Figure S8**.

The diversity of *tox*-positive isolates was evident from their distribution in the *C. diphtheriae* phylogenetic tree, but it was striking that the Gravis branch comprised much less *tox*-positive sublineages than the Mitis branch (**Figure 3**): in the Gravis lineage, there were only three main branches of *tox*-positive isolates: (i) an early-branching group of sublineages; (ii) a branch comprising SL377 and SL384 (two frequent sublineages in France-2022), and (iii) SL8. NTTB isolates were only observed in the Mitis lineage (with one exception in Gravis-SL384) and this phenotype was acquired through multiple independent evolutionary events (**Figure 3**).

A high diversity of *tox*-negative sublineages was also observed in the global dataset: whereas 173 of 245 (70.6%) sublineages were entirely *tox*-negative, only 73 (29.8%) of them had at least 1 *tox*-positive isolate. Of these, 50 sublineages were homogeneous for *tox* status (*i.e*., they included uniquely *tox*-positive genomes), whereas 23 sublineages (9.3%) included both *tox*-positive and *tox*-negative genomes (**Table S1**; **Figure 2**), indicating that the gain or loss of the *tox* gene is not uncommon within sublineages. When considering the genomic clusters, almost all were either *tox*-positive or *tox*-negative in the global dataset. Accordingly, sublineages in the France-2022 dataset were all either *tox* positive or negative, but notably, SL377-GC71 comprised both types of isolates (**Figure 1**).

### 6. Antimicrobial resistance

DIPHTOSCAN includes a screen of *C. diphtheriae* genomes for the presence of antimicrobial resistance genes or mutations against 10 classes of antimicrobial agents. DIPHTOSCAN also computes a resistance score, defined as the number of antimicrobial classes for which at least one resistance gene or mutation is detected. The resistance score varied from 0 to 8 in the global dataset; 38.2% non-redundant global isolates had at least one genomic resistance feature, and 118 isolates (12.1%) were multidrug resistant (acquired resistance to ≥3 drug classes; **Table S1**).

Resistance feature frequencies are shown in **Figure 4B** for the global dataset. The highest frequencies of resistance genes were observed for sulfonamides (exclusively gene *sul1*; rarely present in two copies; 260 non-redundant isolates; 26.6%) and for tetracycline resistance, where *tet(O), tet(W)* and *tet(33)* were present in approximately equal proportions (132 isolates; 13.5% in total). The phenicol resistance gene *cmx* was also commonly found. *pbp2m* was present in 34 (3.5%) isolates, and *ermX* [sometimes named *erm(X)*] in 36 (3.7%) isolates, with 14 (1.4%) isolates carrying both *pbp2m* and *ermX*.

**Figure 4.**
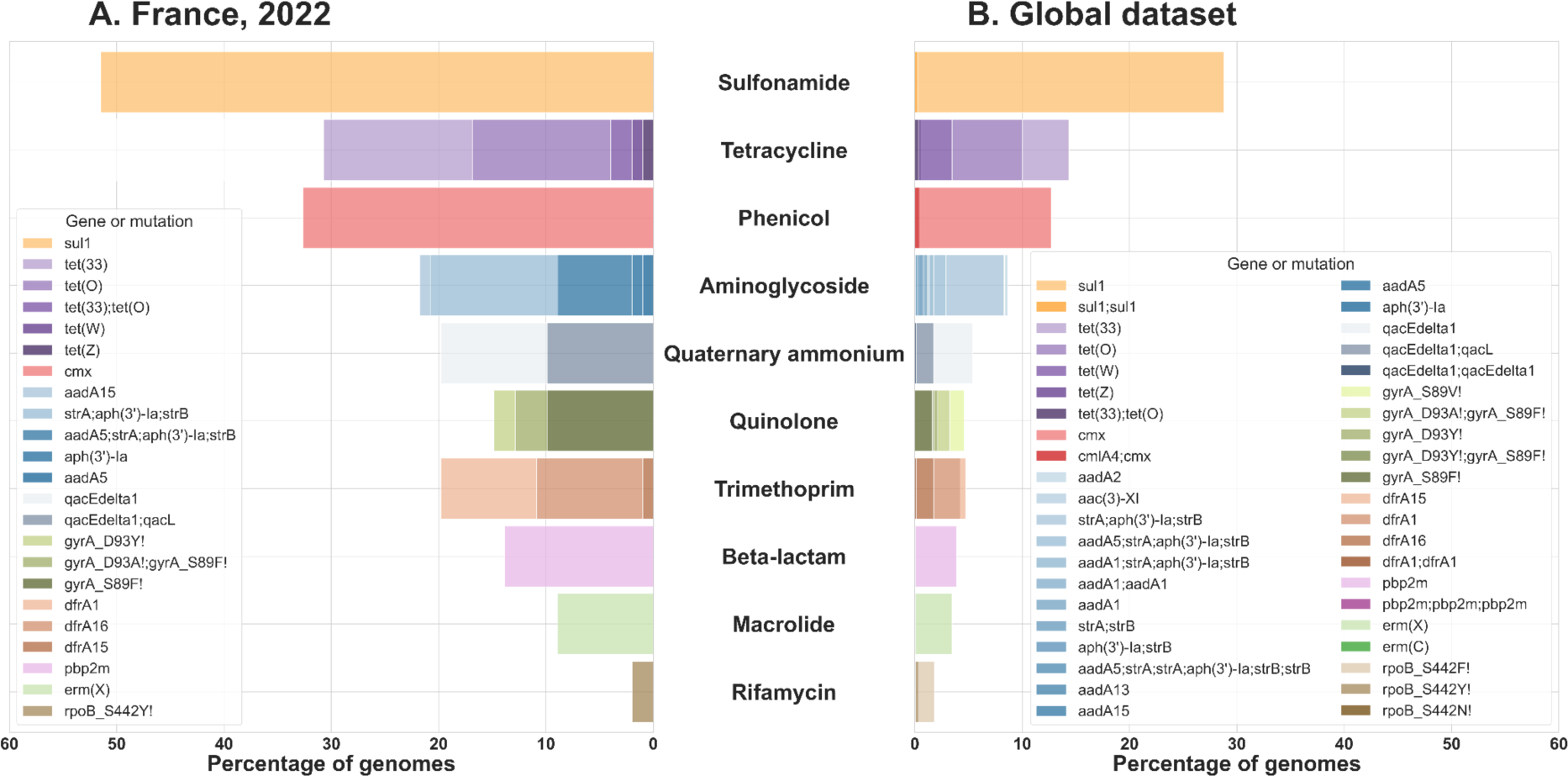
Observed frequencies of resistance genes or mutations. The number of genomes with a genetic feature associated with resistance, per antimicrobial class. Left: Isolates from France, 2022 (n=101 genomes); Right: global deduplicated dataset (n=976 genomes). The bars are ordered vertically by decreasing frequency in the right panel and the bar sectors are colored according to the presence of resistance features (see keys).

Antimicrobial resistance genes were dispersed across the global *C. diphtheriae* phylogenetic tree (**Figure 3**). The distribution of resistance at the sublineage level showed that just above half of the sublineages (128; 52.0%) comprised at least one strain with at least one resistance genomic feature (**Table S1**). The two sublineages with the most resistant strains were SL8 (the main sublineage involved in the ex-USSR outbreak; 46 strains) and SL377 (17 strains) (**Figure 2**). 19 sublineages carried at least one multidrug resistant isolate, and SL377 and SL405 were the most frequent of these (**Figure 2**).

Against this background, the France-2022 isolates appeared to carry resistance features much more frequently, including *pbp2m, ermX* and quinolone-resistance determining mutations (**Figures 1 and 4**). 61 (60.4%) isolates presented at least one resistance feature (**Table S1**; **Figure 1**), and 44 (43.6%) were multidrug resistant.

First-line treatments of diphtheria are penicillin or amoxicillin and macrolides in case of allergy to beta-lactams. The *pbp2m* gene confers decreased susceptibility to penicillin and other beta-lactams (Forde et al., 2020; Hennart et al., 2020), whereas *ermX* (and rarely *ermC*) are associated with erythromycin resistance in *C. diphtheriae* (Tauch et al., 1995, 2003). In the global dataset, 34 isolates (**Table S1;** including strain BQ11 with three copies consistent with Forde *et al*. 2020) carried *pbp2m* and 35 carried *ermX;* 14 (1.4%) isolates carried both genes. Sublineages SL297 and SL484 were the most common carriers of these genes, whereas the frequent multidrug resistant sublineages SL377, SL384 and SL301 did not carry *ermX* and *pbp2m* (**Figure S9**). In France-2022, 8 (7.9%) isolates carried both *pbp2m* and *ermX*. These were observed in patients with travel history from Mali (SL395, SL542, SL852) and Egypt (SL297-GC820).

Antimicrobial susceptibility phenotypes were determined for the France-2022 dataset, and were highly concordant with the presence of resistance features (**Table S4**). Resistance to penicillin and macrolides was associated with *pbp2m* and *ermX*, respectively, although some *ermX*-carrying isolates remained susceptible to erythromycin (**Table S4**).

We included in DIPHTOSCAN a search for integrons, which may harbor multiple resistance genes in *C. diphtheriae* (Barraud et al., 2011; Arcari et al., 2023). In the global dataset, we identified 45 (4.6%) isolates carrying integrons (including integrase-less ones, *i.e*., CALINs) (**Table S1**), which were highly dispersed in the phylogeny (not shown). In France-2022, we found the presence of complete integrons in 9 isolates and integrase-less integrons in 9 additional isolates (18; 17.8%). These structures were strongly associated with antimicrobial resistance, particularly to trimethoprim and sulfonamides (**Figure 1; Table S1**).

### 7. Dual risk isolates: convergence of diphtheria toxin and multidrug resistance, including to first-line treatments

The presence within the same isolates of multidrug resistance and toxigenicity could cause particularly threatening infections. We therefore explored the co-occurrence of these two genotypes (**Figure 2**). In the global dataset, 57 (5.8%) isolates were both multidrug resistant and *tox*-positive. The majority of these isolates belonged to a few sublineages (**Figure 2**), including SL377, which comprised 9 *tox*-positive multidrug resistant isolates mostly from India (and also observed in France-2022). Eight convergent isolates of SL301 were also observed from India, Austria and Syria. SL453 had three *tox*-positive multidrug resistant isolates, which were isolated in Spain and France with links to Afghanistan (Arcari et al., 2023). In metropolitan France, there were 22 *tox*-positive isolates that were multidrug resistant (21.8%), with SL377 and SL696 being predominant among these (**Table S1**, **Figure 1**).

Regarding resistance genes to first-line treatments, there was not a single isolate carrying at the same time *tox, pbp2m* and *ermX* in the global dataset (**Table S1**). However, in France-2022, SL852 isolates (from two patients with travel history from Mali) were *tox*-positive and carried *pbp2m* and *ermX*. Furthermore, they carried other resistance genes including *cmx*, *sul1, dfrA1,* and in addition *tet33* and *aadA15* for isolate FRC1688. This latter isolate only lacked resistance features to quinolones and rifampicin. No other isolate of this particularly concerning sublineage (SL852) was found in the global dataset.

### 8. Lineages Gravis and Mitis differ in the presence of pathogenicity-associated genes

Biovars represent an early attempt to discriminate among *C. diphtheriae* strains (Anderson et al., 1931) and are still commonly reported. We found that lineages Mitis and Gravis, defined genetically based on the presence of the *spuA* gene probably involved in starch utilization, correspond to two distinct parts of the phylogenetic tree (**Figure 3**) as previously reported (Hennart et al., 2020; Guglielmini et al., 2021). Note that the match between lineage and *spuA* or biovar phenotype is not absolute, as a few isolates within the Gravis branch were *spuA*-negative (in particular SL625, SL130, SL102, and SL377) and 42 (5.1%) isolates of the Mitis lineage were *spuA*-positive. Among the France-2022 isolates, for which biovars were in addition determined phenotypically, the two biovars were also phylogenetically distinct (**Figure 1**). Nearly four in five (n=78) of the France-2022 isolates had a Mitis biotype (including 37 *tox*-positives), with 23 Gravis strains (8 *tox*-positive).

To provide a population-level view of pathogenesis features in *C. diphtheriae*, we included in the DIPHTOSCAN database of searched genes, in addition to the *tox* gene, all virulence genes previously demonstrated or strongly suspected to be involved in diphtheria pathogenesis (see **Table S2** for pathogenesis involvement evidence). These include genes involved in iron and heme acquisition, fimbriae biosynthesis and assembly, and other adhesins (Ott et al., 2022).

Screening for these genes in the global dataset revealed highly heterogeneous patterns of presence and phylogenetic distribution (**Table S1**; **Figure S10**). We found that a number of virulence factors are highly conserved within *C. diphtheriae;* for example, DIP1546 was present in all genomes except in 28_DSM43988, and DIP0733, DIP1281, DIP1621, and DIP1880 were fully conserved (**Table S1**). The corynebactin transport (*ciuA-D*) gene cluster was present in all genomes, with one exception, whereas the corynebactin synthesis (*ciuEFG*) locus was absent or incomplete in only 5.4% of genomes (n=29 Mitis, n=25 Gravis); of these, 33 lacked the *ciuE* gene, which is essential for siderophore synthesis. One of the genomes lacking *ciuE* corresponds to the vaccine strain PW8, which is defective for corynebactin synthesis (Russell & Holmes, 1985). The heme-acquisition genes *hmuTUV* were also largely conserved (921 genomes; 94.4%).

In contrast, some genes were infrequent: DIP2014, a gene encoding for a BigA-like adhesin, was detected in only a few sublineages of the Gravis branch (133 isolates), and the DIP0543 (also known as *nanH*, coding for a sialidase) was present in only a few sublineages distributed across the phylogeny (not shown).

Remarkably, we uncovered a sharp divide between lineages Gravis and Mitis in terms of iron metabolism-associated genes, fimbriae gene clusters and other genes (**Figure S10**). The putative siderophore synthesis and transport operon *irp2ABCDEFI-irp2JKLMN* was strongly associated with the Mitis lineage: 513 out of 567 (90.5%) Mitis isolates were *irp2*-positive, whereas only 1 of 406 Gravis isolates was *irp2*-positive. The iron transport cluster *irp1ABCD* was also mainly present in the Mitis lineage. Differently, the *htaA* gene, which is part of the same gene cluster as *hmuTUV* and codes for a membrane protein that binds hemoglobin, was absent or truncated in most genomes from the Mitis branch (92.1%), whereas it was largely conserved in the Gravis branch (99.8% *htaA*-positive). Similar to *htaA*, genes *chtA* and *chtB*, which have sequence and functional similarity to *htaA* and *htaB*, were also strongly associated with the Gravis lineage: 304 of 406 Gravis isolates were *chtAB*-positive (74.9%), whereas only 7 of 567 Mitis isolates were *chtAB*-positive (1.2%). In sharp contrast, the *htaC* gene, which is suspected to be involved in hemin transport, and which is also in genetic linkage with the *hmuTUV* gene cluster, was entirely absent from the Gravis branch, but was detected in 68.6% of Mitis genomes.

Three main fimbriae gene clusters, encoding fimbrial proteins, SpaA, SpaD and SpaH, have been described in *C. diphtheriae* (Rogers et al., 2011; Reardon-Robinson & Ton-That, 2014; Sangal & Hoskisson, 2016). We found that these were more commonly found in the Gravis branch compared to the Mitis branch (**Figure S10**). The SpaH gene cluster (*spaGHI-srtDE*) was present in its entirety in 254 genomes and as a cluster with one missing gene in 29 isolates, all of which belonged to the Gravis lineage. The other two systems showed some variability in the distribution of their genes. The sortase-mediated assembly genes of the SpaA type pili, *spaABC,* were found in biovar Gravis in similar proportions (87.2% *spaA*, 86.2% *spaB* and 86.0% *spaC*-positive), whereas in Mitis *spaB* was present in about half of the genomes (49.0%) and *spaA* and *spaC* in one third (17.5%, and 18.2%, respectively). The distribution of the SpaA pilin-specific sortase gene *srtA* was similar to that of *spaB* (98.8% in Gravis, 49.9% in Mitis), and the complete SpaA gene cluster *spaABC-srtA* was found in only 299 genomes (30.6%), the majority of which were of Gravis lineage (n=256). Last, genes of the SpaD cluster were less frequent (*spaD* 8.7%, *spaE* 14.9%, *spaF* 9.3%, *srtB* 33.2%, *srtC* 33.7%) compared to the other pili types, and the complete gene cluster (*spaDEF-srtBC*) was found only in 11 genomes, all of which belonged to lineage Gravis. Interestingly, the presence of SpaD and SpaH complemented each other in the Gravis branch (**Figure S10**).

We further found that the collagen-binding protein DIP2093 (Peixoto et al.,2017) is strongly associated with the Gravis lineage: 118 of 406 (29.1%) Gravis isolates were DIP2093-positive, whereas only 3 of 567 (0.5%) Mitis isolates were.

The complement of virulence genes of the France-2022 isolates was in full agreement with their Gravis/Mitis placement and the above observations. For example, the *irp2A-I* and *irp2J-N* gene clusters were present uniquely in sublineages belonging to the Mitis branch, and the *htaC* gene was present only in 64.2% of the Mitis genomes (**Table S1**); *chtA* and *chtB* were completely absent in Mitis and the collagen-binding protein DIP2093 uniquely in Gravis isolates (n=16, 47.1%). None of the France-2022 isolates carried a complete SpaD fimbriae cluster; in particular, they all lacked at least the *spaD* gene; and only 8 Gravis genomes carried the complete SpaH cluster. The latter were dispersed among various lineages (SL32, SL374, SL502, SL542, SL130).

## Discussion

In recent years, large epidemics of diphtheria have been observed, *e.g.,* in South Africa, Bangladesh and Yemen (du Plessis et al., 2017; Polonsky et al., 2021; Badell et al., 2021), while a progressive increase of diphtheria cases has been noted in multiple countries (Bernard et al., 2019; Truelove et al., 2020). However, so far, our understanding of diphtheria reemergence has been hindered by a lack of background knowledge on the population diversity of *C. diphtheriae*, its sublineages of concern and the epidemiology of their local or global dissemination. Here, we report on a sharp increase in *tox*-positive *C. diphtheriae* in France in 2022, and developed a bioinformatics pipeline, DIPHTOSCAN, which enables to harmonize the way genomic diversity and genetic features of medical concern are detected, reported and interpreted. We illustrate how this novel tool provides clinically-relevant genomic profiling and evolutionary understanding of emergence, by placing the 2022 *C. diphtheriae* from France in the context of 1,249 global *C. diphtheriae* genomes.

Our results provide an updated overview of the population diversity of *C. diphtheriae* based on currently available genomic sequences. As previously reported (Berger et al., 2019; Seth-Smith & Egli, 2019; Hennart et al., 2020; Guglielmini et al., 2021), *C. diphtheriae* is made up of multiple sublineages that are related through a star-like phylogeny. We here uncovered 123 novel sublineages, for a total of 253 described ones. We observed that, compared to previous datasets, there was no sublineage fusion upon adding novel genomes, which indicated an excellent stability of *C. diphtheriae* sublineage classification. The latter provides a broad classification of isolates that correlates strongly with classical MLST, and which facilitates a deep-level approach to *C. diphtheriae* diversity and evolution. The naming of sublineages by inheritance of ST numbers will facilitate continuity with classical MLST. Besides, sublineage classification is more congruent with phylogenetic relationships: whereas most (140/146; 95.8%) non-singleton sublineages were monophyletic, only 134 of 167 (79.8%) non-singleton STs were (data not shown). We therefore strongly recommend transitioning from MLST to the cgMLST-based nomenclature, which is available on the BIGSdb-Pasteur platform. Our phylogenetic analysis of reference strains of the historical ribotype nomenclature provides a first overview of their relationships, to our knowledge, and allows revisiting genealogical inferences that were made among ribotypes based on CRISPR spacer variation (Mokrousov, 2009).

Genomic clusters represent a much narrower genetic classification of *C. diphtheriae* isolates, compatible with recent transmission (Guglielmini et al., 2021). Therefore, genomic clusters appear more relevant than sublineages for epidemiological investigation purposes, as illustrated for example within SL377: whereas GC817 was associated with Afghanistan, GC71 was associated with Senegal and these two genomic clusters of sublineage SL377 were clearly distinct phylogenetically (**Figure 1**).

The diagnostic and surveillance of diphtheria is largely based on the detection of the *tox* gene and its expression (WHO, 2018). We found that the determination of the *tox* gene presence by DIPHTOSCAN was highly concordant with the experimental reference qPCR. We also found that DIPHTOSCAN can predict a large proportion of non-toxigenic *tox* gene-bearing (NTTB) isolates. Still, some NTTB isolates were not identified by DIPHTOSCAN. These cases may be attributable to (i) a lack of detection by the Elek test due to a low level of expression of the toxin gene in some strains, or (ii) yet unknown genetic mechanisms that abort *tox* gene expression entirely (unexplained true NTTB). Future work is needed to define the genotype-phenotype links underlying toxigenicity and to improve our predictive capacity of toxigenicity from genomic sequences. In the non-redundant global dataset, 16.0% of *tox*-positive isolates were predicted as NTTB, which provides a quantitative view of the relevance of differentiating mere *tox* gene presence from actual toxigenicity. The capacity to predict toxigenicity from sequences opens interesting perspectives as to the diagnostic of diphtheria based on rapid genomic sequencing. Our phylogenetic analysis showed that gain or loss of the *tox* gene is a rare event at the timescale of genomic cluster diversification. The phenomenon of *tox* status switch by phage acquisition or loss during infection or transmission was suspected previously (Pappenheimer & Murphy, 1983) and deserves further study given its importance for public health and clinical management.

Up until now, antimicrobial resistance has been considered of moderate clinical concern in *C. diphtheriae* (Zasada, 2014; WHO, 2018). Although resistant strains have been described, clinical susceptibility breakpoints have lacked standardization and the prevalence, origin and dissemination of resistance genetic features are largely unknown. Here, we identified in the France-2022 isolates as well as in the global *C. diphtheriae*, multidrug resistant isolates and/or isolates resistant to first-line treatments. We provide an overview of the prevalence and distribution of resistance genes or mutations in *C. diphtheriae*, and identify sublineages that carry multiple resistance genes. Because antimicrobial resistance phenotypes are typically unattached to publicly available genomic sequences, it is not possible to link these genomic features complements to resistance phenotypes systematically. However, this (**Table S4**) and previous works clearly showed that most resistance genetic features identified here may impact resistance phenotypes (Tauch et al., 1995, 2003; Hennart et al., 2020; Forde et al., 2020). Of particular concern, *tox*-positive isolates that are resistant to multiple drugs and/or first-line treatments were identified herein, with the convergence of *tox*, *pbp2m* an *ermX* in two 2022 cases with a travel history from Mali, which were resistant to 9 and 11 out of 23 tested antimicrobials, respectively. Such isolates may pose serious clinical management difficulties, and multidrug resistant *C. diphtheriae* should therefore be closely monitored.

The combined analysis of the France-2022 and global datasets using a unique pipeline provides context to the reemergence of diphtheria (**Figure S6**). The occurrence of cases of diphtheria among migrants, the vast majority of whom are not up to date with their vaccinations, raises concerns of the emergence of cluster cases in accommodation facilities for migrants, refugees or asylum seekers (Badenschier et al., 2022; Kofler et al., 2022). Professionals dealing with these populations need to be particularly vigilant in spotting clinical signs of diphtheria and ensuring that their vaccinations are up to date. Here, we found that some sublineages contributing to the reemergence were previously observed, whereas others are described for the first time. For example, SL377, one of the major toxigenic and resistant sublineages observed in France-2022, had been circulating in India during 2016 and was reported in Europe (Spain and France) since 2015 (**Table S1**). In contrast, SL698 was absent from the global dataset. Of the 10 *tox*-positive France-2022 sublineages, five were associated with travel from Afghanistan, and were recently described in other European countries too (Badenschier et al., 2022; Kofler et al., 2022).

The DIPHTOSCAN tool will facilitate the harmonized characterization of *C. diphtheriae* sublineages of concern. Several virulence-associated genes were largely conserved in the entire *C. diphtheriae* population analyzed; these genomic features may therefore be central for *C. diphtheriae* colonization and transmission among humans, as there appears to be a strong selective pressure to maintain them. The distribution of other, more variably present, virulence-associated genes uncovers a very striking dichotomy between the Gravis and Mitis lineages, as heme and iron-acquisition systems and Spa-encoded fimbriae gene clusters were either associated with the Mitis or the Gravis lineages, in a largely mutually exclusive way. Based on these observations, the Gravis lineage may preferentially capture iron from hemin, whereas the Mitis one could be associated with the ability to synthesize and use siderophores. There might be important implications for the regulation and expression level of the *tox* gene, which is controlled by the iron-dependent DtxR repressor. Importantly, the toxin gene and its NTTB-leading disruptions were also unequally distributed between Gravis and Mitis lineages. It was noted early that toxin production is less inhibited by infection-relevant iron concentrations in Gravis strains (Mueller, 1941; McLeod, 1943), and our results shed a new light and provides experimentally testable hypotheses on this critical difference in the biology of infection of the Gravis and Mitis lineages.

Another striking feature we uncovered is the distribution of gene clusters coding for fimbriae. Previous work reported SpaA as being largely conserved in *C. diphtheriae*, with SpaD and SpaH being more variably present (Reardon-Robinson & Ton-That, 2014; Sangal & Hoskisson, 2016; Ott, 2018). We found that SpaA was largely present in our dataset, however, the complete gene cluster *spaABC-srtA* was mostly found in the Gravis branch. SpaD was also more common among Gravis genomes, although the complete cluster (*spaDEF-srtBC*) was only detected in a minority of genomes. None of the Mitis isolates were positive for SpaH. These three Spa systems were experimentally shown to be involved in adhesion to different human tissues: pharyngeal (SpaA), laryngeal (SpaD) and pulmonary (SpaH) epithelial cells (Mandlik et al., 2007; Reardon-Robinson & Ton-That, 2014). The Gravis/Mitis dichotomy in Spa-type fimbriae may have important implications regarding a possible differential ecology, transmission, tissue tropism and pathophysiology of these two major *C. diphtheriae* lineages.

In conclusion, we developed and applied to a large dataset, the bioinformatics tool DIPHTOSCAN. Its public availability and ease of use will enable to conveniently extract and interpret genomic features that are relevant to the clinical and public health management of diphtheria cases, to understand the microbiological determinants of (re)emerging sublineages, and to future research on the genotype-clinical phenotype links in *C. diphtheriae*. This dedicated tool is also applicable to the other members of the *C. diphtheriae* complex, such as *C. ulcerans* (data not shown). Harmonization of genomic studies in this group of pathogens, which have been largely forgotten but currently undergo re-emergence in Europe and elsewhere, will support genomic surveillance of diphtheria, will contribute to enhance our understanding of the pathogenesis of modern diphtheria, and opens interesting hypotheses as to the underlying mechanisms of variation in clinical severity and forms of diphtheria.

## Acknowledgements

We thank Martin Maiden and Keith Jolley (Oxford University) for maintaining the previous MLST data from Oxford’s PubMLST database and for providing the data for import into the BIGSdb-Pasteur *C. diphtheriae* species complex database.

## Author contributions

S. Brisse (S.B.) conceived, designed, and coordinated the study. Melanie Hennart (M.H.) developed the DIPHTOSCAN tool with input from SB. M.H. and S.B. analyzed the genomic data. M.H. created the figures and tables. S.B. and M.H. created the first draft of the manuscript, worked together to improve it and reviewed the final version. Chiara Crestani analyzed the iron metabolism and fimbriae genes distribution and wrote the first version of the corresponding sections. Sebastien Bridel performed the merger of the Oxford PubMLST and BIGSdb-Pasteur databases. Annick Carmi-Leroy, Sylvie Brémont, Annie Landier, Nathalie Armatys and Virginie Passet provided technical assistance with the microbiological characterization and sequencing of the *C. diphtheriae* isolates. Edgar Badell and Julie Toubiana contributed to the NRC operations coordination. Laure Fonteneau and Sophie Vaux coordinated diphtheria epidemiological surveillance in France. All authors reviewed and approved the final contents of the manuscript.

## Data, scripts, code, and supplementary information availability

The latest version of the DIPHTOSCAN code will be available at https://gitlab.pasteur.fr/BEBP/diphtoscan and the version used in this work in available at: https://zenodo.org/record/7774709.

The genome sequence data generated in this work has been made publicly available through NCBI/ENA bioproject PRJEB22103 (https://www.ebi.ac.uk/ena/browser/view/PRJEB22103).

The trees are available at https://itol.embl.de/shared/Pasteur_BEBP in the projet: ‘Hennart et al., 2023: diphtOscan’.

The supplementary appendix is available in zenodo at: https://doi.org/10.5281/zenodo.8123234

### Ethical approval statement

Diphtheria is a notifiable disease in France. Phenotypic and genotypic analyses of bacterial isolates were carried out within the framework of the mandate given to the National Reference Center for Corynebacteria of the Diphtheriae Complex by the Ministry of Health (Public Health France). All French bacteriological samples and data were collected in the frame of the French national diphtheria surveillance and are collected, coded, shipped, managed and analyzed according to the French National Reference Center protocols. Other strains were obtained from culture collections.

### Authors’ license statement

This research was funded, in whole or in part, by Institut Pasteur and by European Union’s Horizon 2020 research and innovation programme. For the purpose of open access, the authors have applied a CC-BY public copyright license to any Author Manuscript version arising from this submission.

