## Supplementary appendix for "A global *Corynebacterium diphtheriae* genomic framework sheds light on current diphtheria reemergence"

**Contents**:

Supplementary tables S1 to S5

Supplementary figures S1 to S10

### Supplementary tables

**Table S1.** Characteristics of the 1,350 *Corynebacterium diphtheriae* isolates

**Table S2.** Virulence-associated genes

**Table S3.** Zone diameter interpretative breakpoints used for antimicrobial agents

**Table S4.** Antimicrobial susceptibility phenotypes of the France-2022 isolates

**Table S5.** Count data for all figures

- *See separately attached tables -*

#### Supplementary figures

**Figure S1.** Epidemiological curve of *tox*-positive *C. diphtheriae* infections in France

**Figure S2.** Flowchart of the main analytical steps of DIPHTOSCAN

**Figure S3.** Bioinformatics tools integrated into DIPHTOSCAN

**Figure S4.** Custom features of DIPHTOSCAN database

**Figure S5.** Sublineage frequency comparisons between datasets

**Figure S6**. Phylogenetic tree of 1,350 *Corynebacterium diphtheriae*

**Figure S7**. Phylogenetic position of the ribotype reference strains of *Corynebacterium diphtheriae*

**Figure S8.** Phylogenetic tree of amino acid-translated *tox* alleles found in the 1350 genomes dataset

**Figure S9**. Sublineage distribution of resistance features, including *pbp2m* and *ermX*

**Figure S10**. Phylogenetic distribution of iron metabolism and adhesion-associated genes in *Corynebacterium diphtheriae* lineages Mitis and Gravis

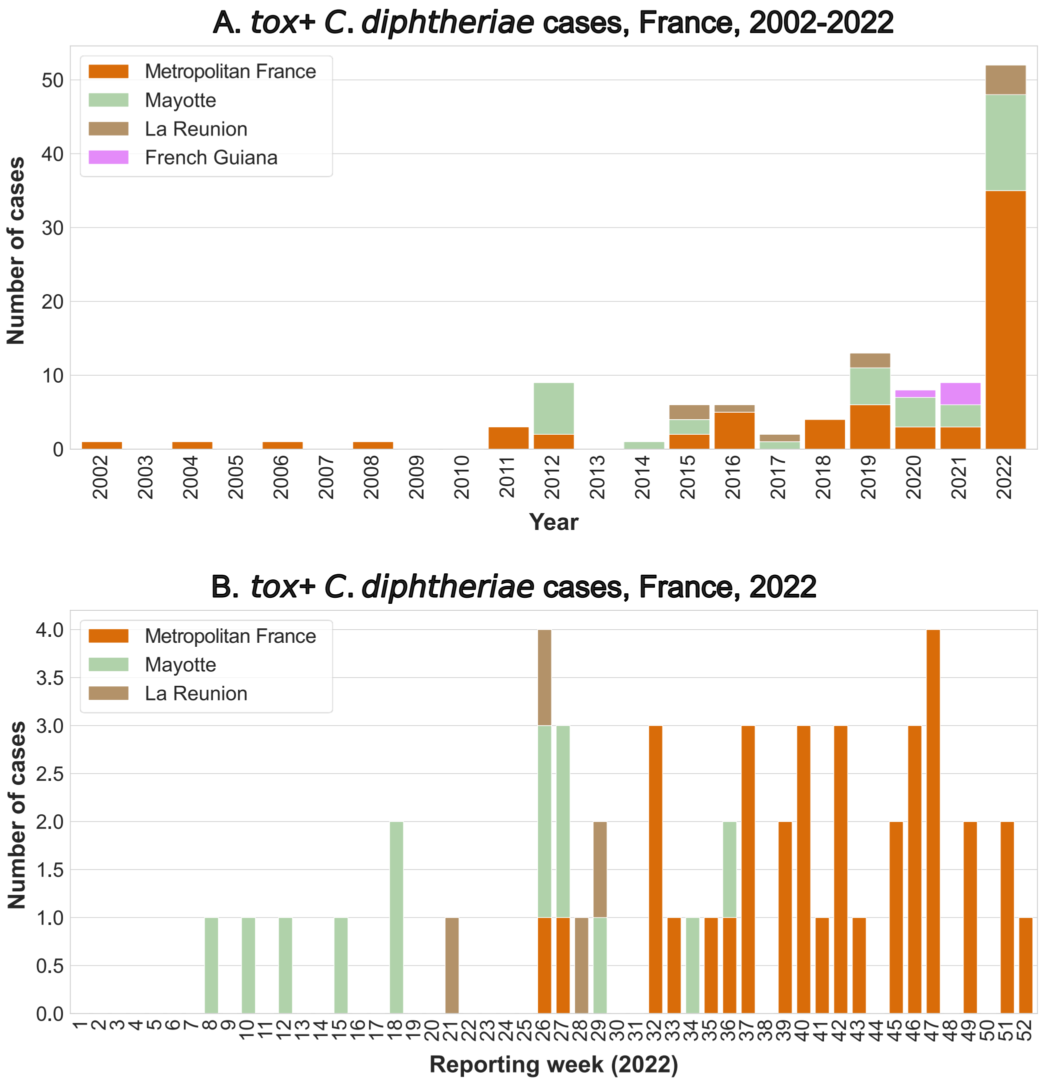


**Figure S1. Epidemiological curve of *tox*-positive *C. diphtheriae* infections in France**

**A**. Number of reported cases by year, France, 2002-2022; **B**. Number of reported cases by week, France, 2022. The cases were either from metropolitan France, or overseas territories (see color key).

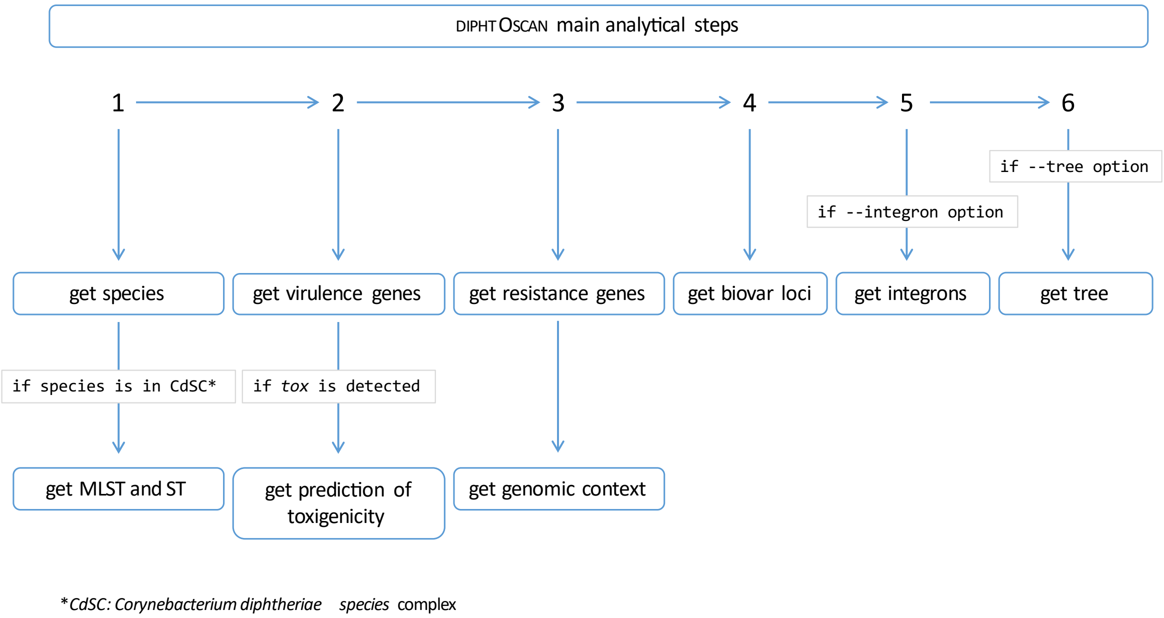


**Figure S2. Flowchart of the main analytical steps of DIPHTOSCAN**

DIPHTOSCAN starts with a taxonomic check, and then proceeds to virulence genes detection (including the tox gene, and its possible truncations), resistance features (genes and mutations, and their co-localization), and biovar-associated genes (*spuA* and *nar* clusters). Optionally, the presence of integrons is detected, and a k-mer distance-based tree can be built.

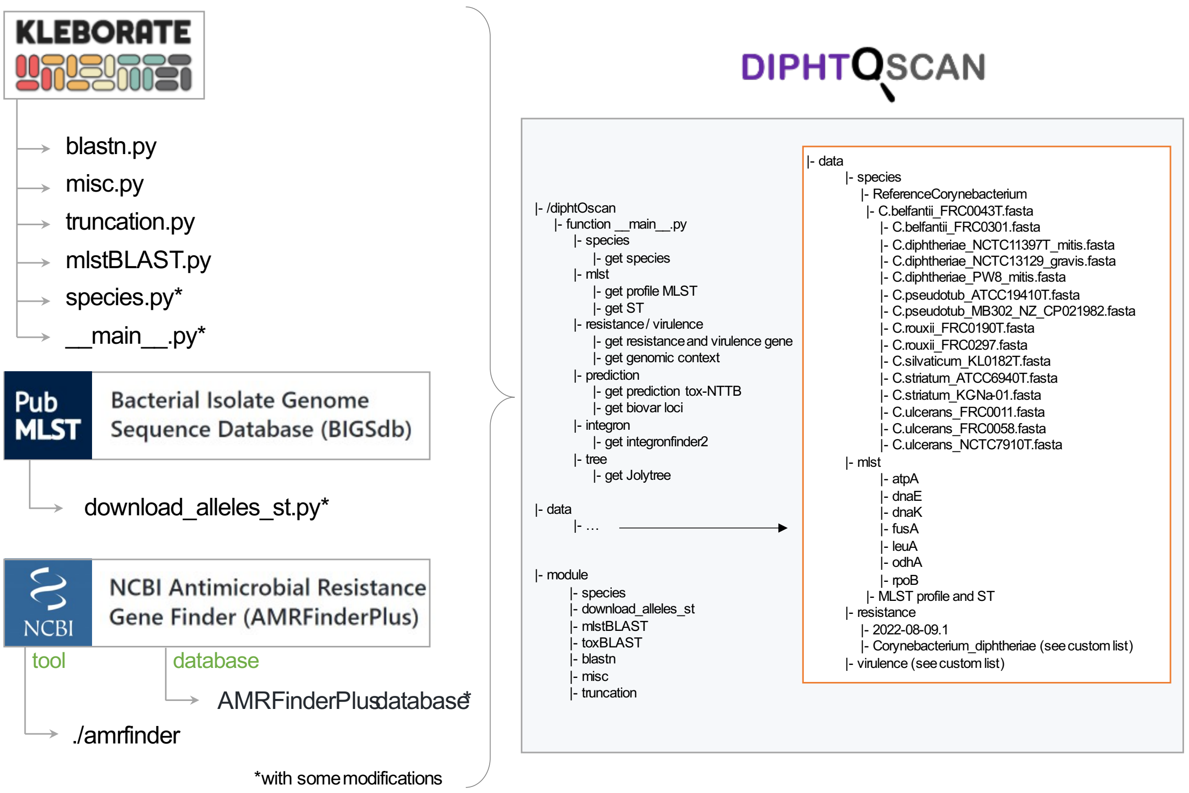


**Figure S3. Bioinformatics tools integrated into DIPHTOSCAN**

DIPHTOSCAN is a pipeline which groups a set of novel scripts and adaptations of previous tools, including Kleborate, BIGSdb, AMRFinderPlus and IntegronFinder2.

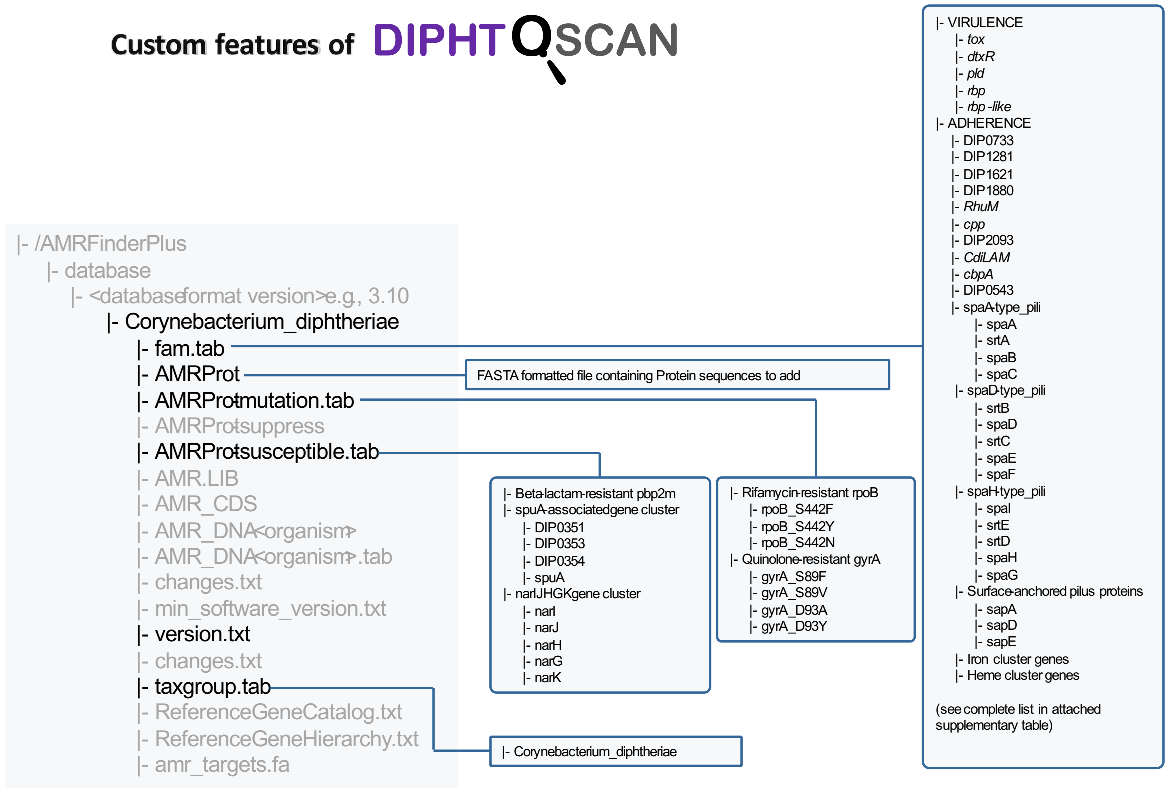


**Figure S4. Custom features of DIPHTOSCAN database**

Description of the custom features of the DIPHTOSCAN database. The database was built by adding to the AMRFinderPlus database, *ad-hoc* resistance and virulence genes, as well as molecular determinants to predict the biovar.

#### A. Effect of deduplication of the global dataset

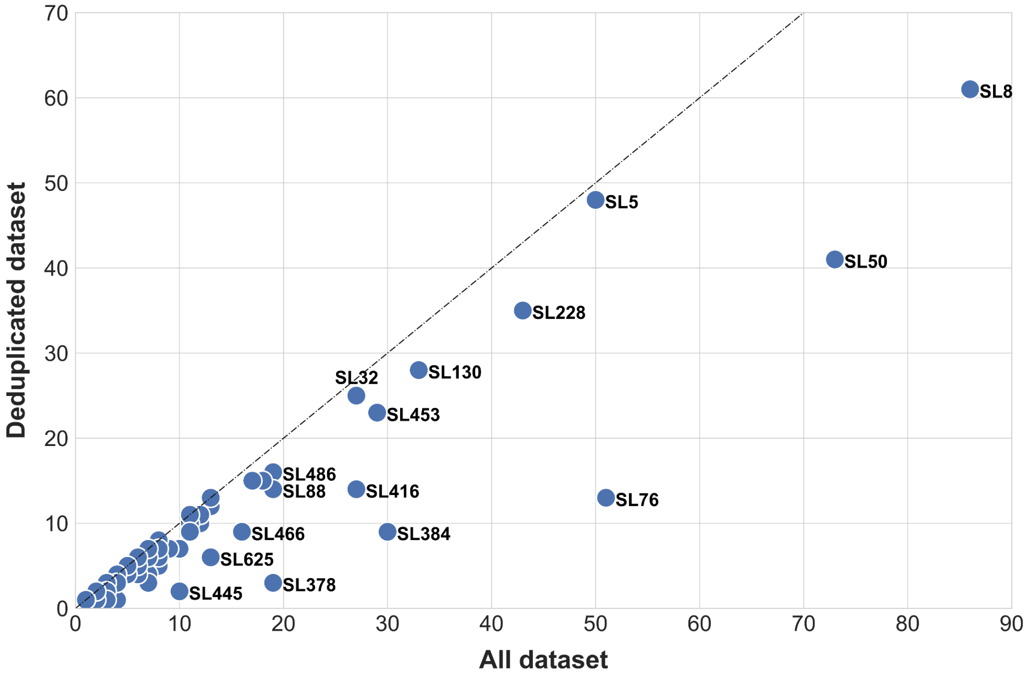


#### B. Sublineage representation in France-2022 versus the deduplicated dataset

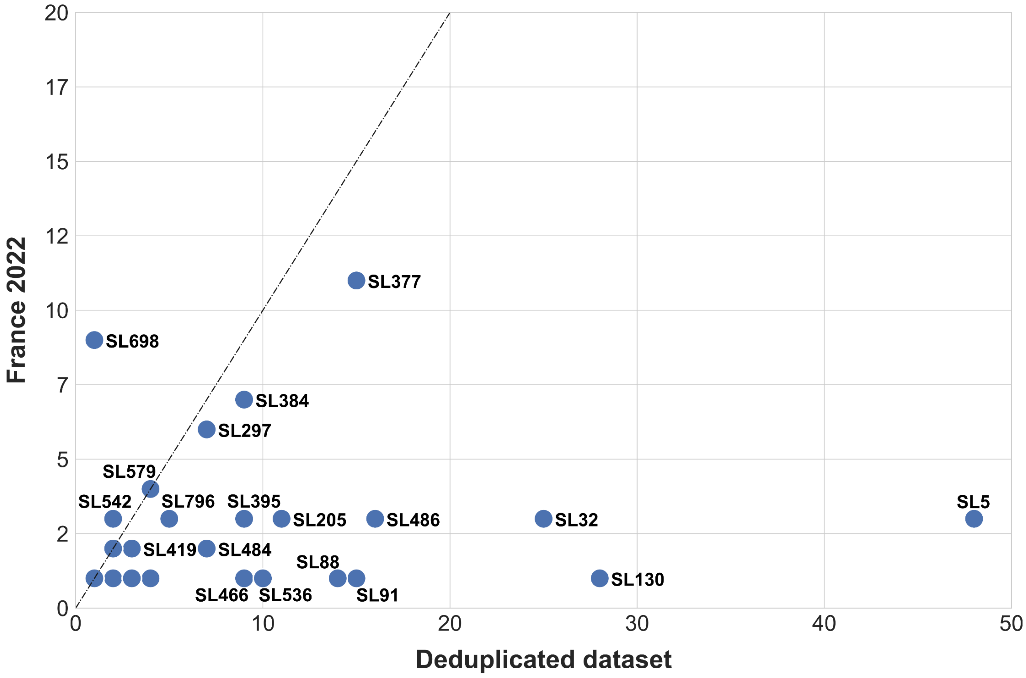


**Figure S5. Sublineage frequency comparisons between datasets**

**A.** Effect of deduplication of the global dataset. X-axis: The number of isolates in the entire global dataset (n=1,249 genomes). Y-axis: The number of isolates in the deduplicated global dataset (n=976). Each sublineage is represented as a circle; the main sublineages' identifiers are shown.

**B**. Sublineage representation in France-2022 *versus* the global (deduplicated) dataset. X-axis: The number of isolates in the deduplicated dataset (n=976 genomes). Y-axis: The number of isolates in the France-2022 dataset (n=101). Each sublineage is represented as a circle; the main sublineages' identifiers are shown.

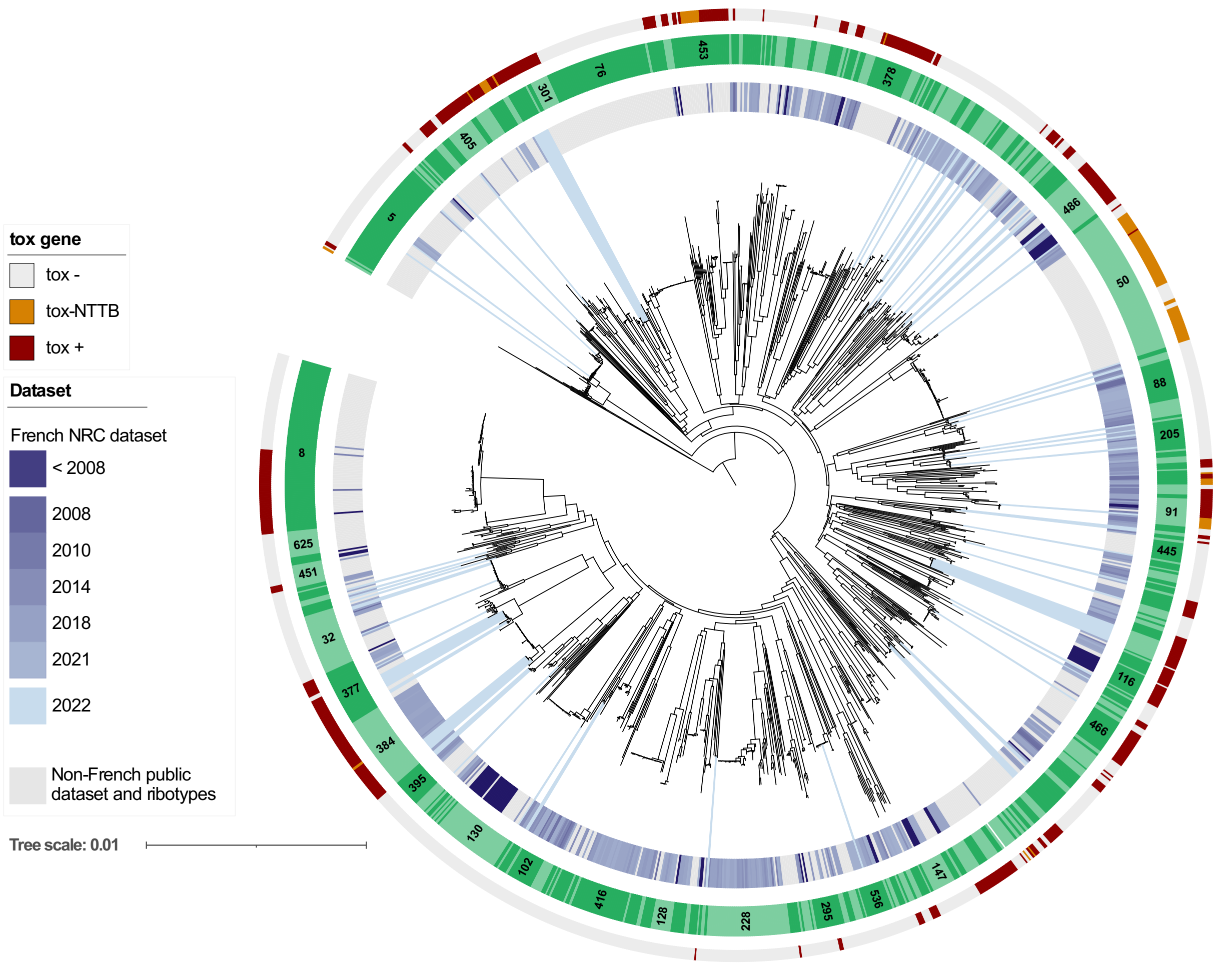


**Figure S6. Phylogenetic tree of 1,350 *Corynebacterium diphtheriae***

The tree shows the placement of the France-2022 isolates (n=101) within the diversity of the 1,249 global non-deduplicated genomes. The tree was generated using JolyTree (Criscuolo, 2020). The scale bar represents the number of nucleotide substitutions per site. The first circle indicates the dataset, with France-2022 isolates indicated with light-blue rays from the tree leaves, and a blue color gradient (darker blue for older isolates) for French NRC isolates prior to 2022. Grey color sectors correspond to the global genomes. The second inner circle indicates sublineage alternation; main sublineages are labeled within the sectors. The following circle indicates the presence, disruption or absence of the diphtheria toxin *tox* gene (see key).

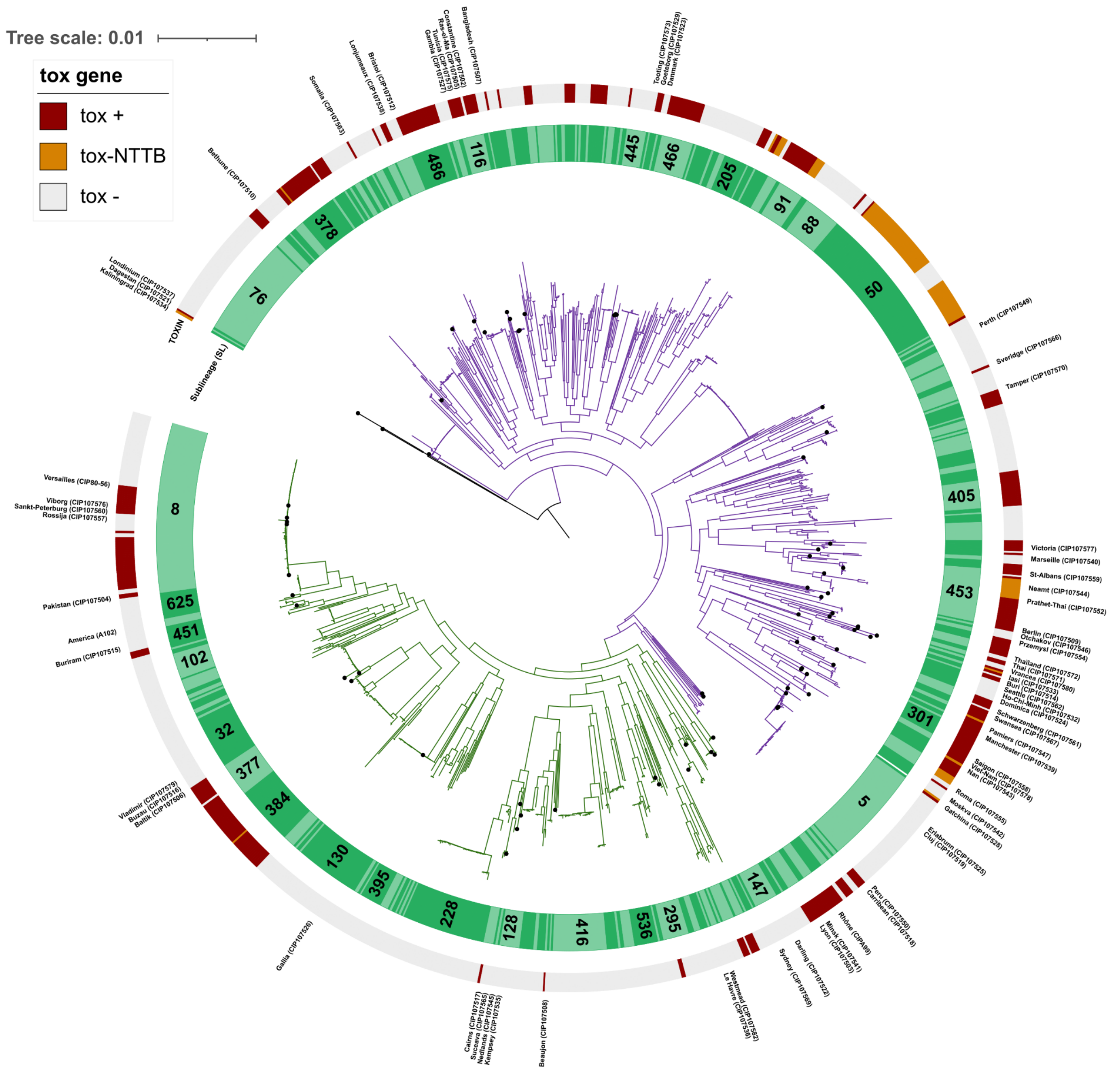


**Figure S7. Phylogenetic position of the ribotype reference strains of *Corynebacterium diphtheriae***

The tree was obtained by maximum likelihood based on a multiple sequence alignment of the core genome, and was rooted with *C. belfantii* (not shown; same tree as in main Figure). The scale bar gives the number of nucleotide substitutions per site. The inner circle indicates sublineage alternation; main sublineages are labeled within the sectors. The second circle indicates the presence, disruption or absence of the diphtheria toxin *tox* gene (see key). Each point on the tree leaves corresponds to a ribotype reference strain. The ribotype names and reference strain identifier are indicated outside the circles.

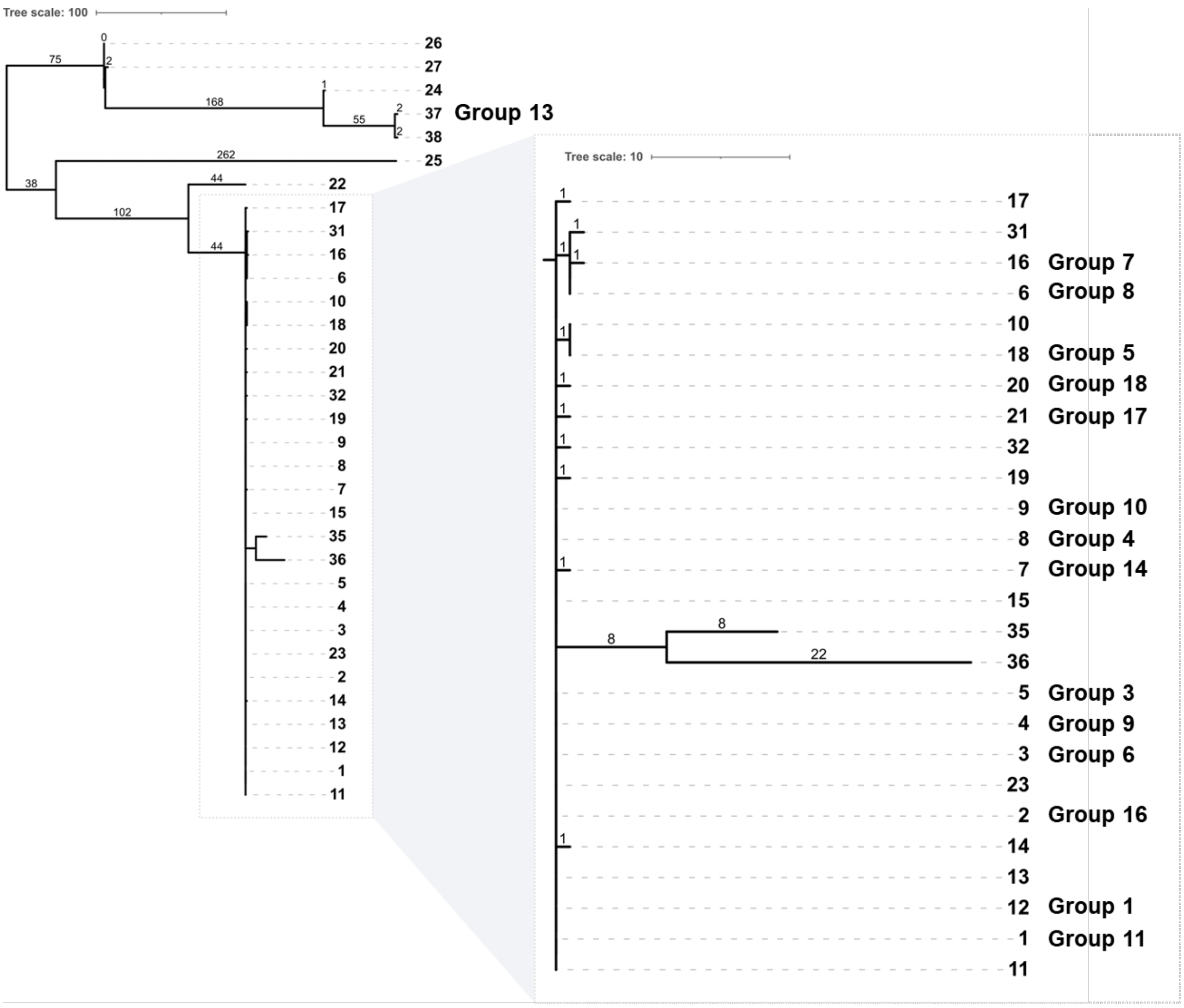


**Figure S8. Phylogenetic tree of amino acid-translated *tox* alleles found in the 1350 genomes dataset.**

The Neighbor-Joining tree was built based on amino acid-translated *tox* alleles. The scale bars give the number of substitutions, as indicated above the branches. These leaves are labeled with integer numbers, which correspond to the *tox* alleles from the BIGSdb nomenclature (https://bigsdb.pasteur.fr/cgi-bin/bigsdb/bigsdb.pl?db=pubmlst_diphtheria_seqdef&page=alleleQuery&locus=tox&submit=1); and the groups indicated after the leaves are those given in the study by Will *et al.*, 2021 (DOI: 10.1038/s41467-021-21870-5). The correspondence between the *tox* alleles of this study and that of Will *et al.*, 2021, was achieved by using the common isolates available in the genomic dataset; *tox* alleles with no correspondence shown here, were not present among the isolates studied in Will *et al.*

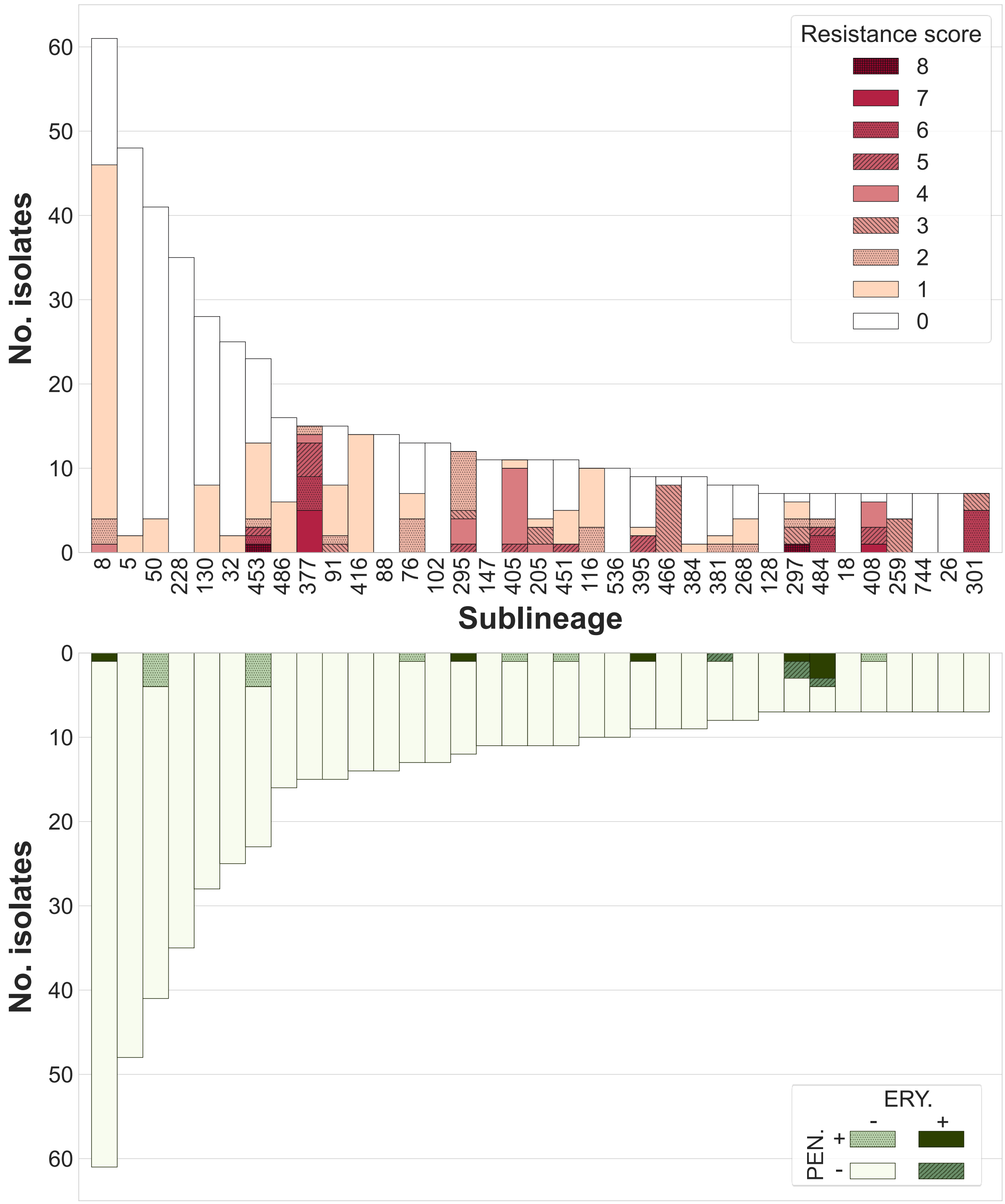


**Figure S9. Sublineage distribution of resistance features, including *pbp2m* and *ermX***

(Top) The bars height corresponds to the number of genomes per sublineage (entire dataset, 1,249 isolates). Bars are colored according to the resistance score, i.e. the number de resistance families present in the strains (see color key).

(Bottom) Same as in A, with genomes carrying genes associated with resistance to penicillin (PEN.) and/or erythromycin (ERY.) resistance being colored (see key).

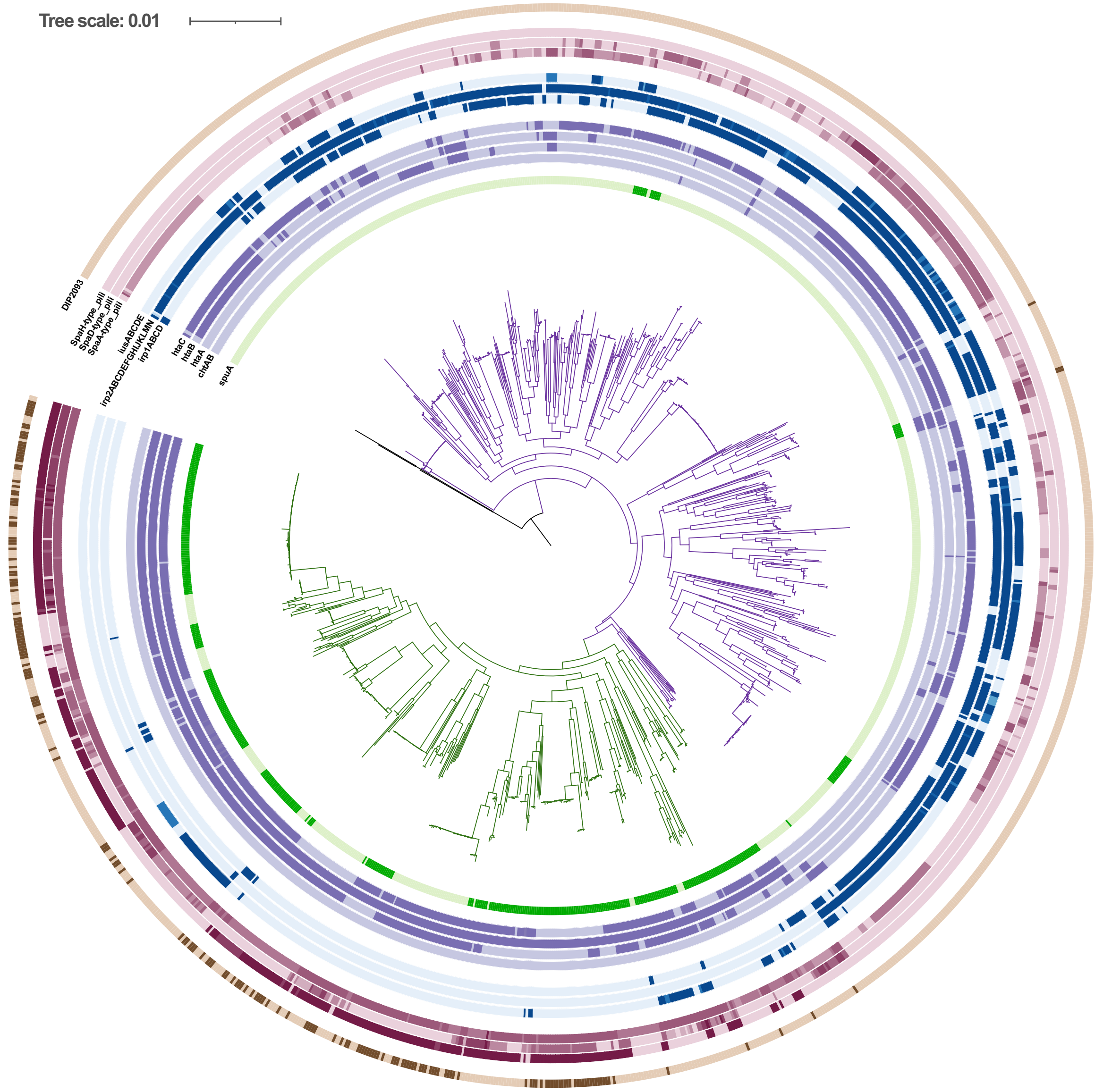


**Figure S10**. **Phylogenetic distribution of iron metabolism and adhesion-associated genes in *Corynebacterium diphtheriae* lineages Mitis and Gravis**

The tree was obtained by maximum likelihood based on a multiple sequence alignment of the core genome (as in main Figure). The scale bar gives the number of nucleotide substitutions per site. The main lineages Mitis and Gravis are drawn using purple and green branches, respectively. The first inner circle indicates the presence of the *spuA* gene (darker color: presence). The four following circles indicate the presence of the heme metabolism-associated genes *chtAB*, *htaA*, *htaB* and *htaC*. The three following circles indicate the presence of the iron metabolism-associated genes: *irp1ABCD*, *irp2ABCDEFI-irp2JKLMN* and *iusABCDE*. Next, three following circles indicate the presence of the fimbriae gene clusters: SpaA-type, SpaD-type and SpaH-type. The most external circle indicates the presence of collagen-binding protein DIP2093. For circles that represent clusters of genes, a color gradient represents the number of genes present in the cluster (darker: more genes).
